# Convert Your Favorite Protein Modeling Program Into A Mutation Predictor: “MODICT”

**DOI:** 10.1101/038992

**Authors:** Ibrahim Tanyalcin, Katrien Stouffs, Dorien Daneels, Carla Al Assaf, Willy Lissens, Anna Jansen, Alexander Gheldof

**Affiliations:** Center for Medical Genetics, UZ Brussel, Laarbeeklaan 101, 1090 Brussel, Belgium.; Neurogenetics Research Group, Reproduction Genetics and Regenerative Medicine Research Group, Vrije Universiteit Brussel(VUB), Laarbeeklaan 101, 1090 Brussel, Belgium.; Center for Medical Genetics, Reproduction and Genetics, Reproduction Genetics and Regenerative Medicine, Vrije Universiteit Brussel(VUB), UZ Brussel, Laarbeeklaan 101, 1090 Brussel, Belgium.; Pediatric Neurology Unit, Department of Pediatrics, UZ Brussel, Laarbeeklaan 101, 1090 Brussel, Belgium.; Center for Human Genetics, KU Leuven and University Hospitals Leuven, Herestraat 49, 3000 Leuven, Belgium.

**Keywords:** prediction, 3D protein model, bioinformatics

## Abstract

**Motivation:** Predict whether a mutation is deleterious based on the custom 3D model of a protein.

**Methods:** We have developed MGDIGT, a mutation prediction tool which isbased on per residue rmsd (root mean square deviation) values of superimposed3D protein models. Our mathematical algorithm was tested for 42 described mutations in multiple genes including renin, beta-tubulin, biotinidase,sphingomyelin phosphodiesterase-1, phenylalanine hydroxylase and medium chainAcyl-Coa dehydrogenase. Moreover, modict scores corresponded toexperimentally verified residual enzyme activities in mutated biotinidase,phenylalanine hydroxylase and medium chain Acyl-CoA dehydrogenase. Several commercially available prediction algorithms were tested and results werecompared. The modict PERL package and the manual can be downloaded from https://github.com/MODICT/MODICT.

**Conclusion:** We show here that modict is capable tool for mutation effect prediction at the protein level, using superimposed 3D protein models instead of sequence based algorithms used by PGLYPHEN and SIFT.

## 1 Introduction

### 1.1 State of the art

As next generation sequencing (NGS) is advancing the field of molecular biology today, more human protein variants are identified than ever before. One of the greatest challenges in this field is to be able to predict whether the detected variants are real disease-causing changes underlying the patients condition.

The current concept of mutation effect prediction heavily depends on the composite algorithms that mainly implement a sequence-based BLAST search that tries to identify a number of similar protein sequences above a preset threshold, then relate and combine several other parameters such as PSIC (Position-Specific Independent Counts), known three-dimensional (3D) structures of similar proteins, surface area, *β*-factor and atomic contacts. Some available algorithms (e.g.PolyPhen 2, http://genetics.bwh.harvard.edu/pph2/, [1]) use all above whereas others use either a portion or a more diverse set of parameters (e.g.SIFT (http://sift.jcvi.org/, [2]), MUTATION TASTER (http://www.mutationtaster.org/, [3]), PROVEAN (http://provean.jcvi.org/index), [4]). Nonetheless, the fact that these algorithms take into account non-mutually exclusive (non-orthogonal) features, the method to correctly combine the results to derive a conclusive output remains ambiguous. One recently described method uses weighted means obtained from false positive rates and false negative rates of each distinct algorithm to approach a consensus score (Condel: http://bg.upf.edu/condel/home [5]). Even after utilizing cancer-trained methods, such integration of scores were not able to correctly classify all variants [6].

### 1.2 Hypothesis and problem definition

A high percentage of genomic variants in protein-coding genes were shown to modify the tertiary structure of the coded protein sequence. These structural modifications can be predicted by comparing the 3D structures of the wild type and mutant protein (.pdb files). The 3D structures are generated in commercial or academic-only servers and software (l-TASSER, http://zhanglab.ccmb.med.umich.edu/I-TASSER/ [7,8], SWiSS-MODEL http://swissmodel.expasy.org/ [9], MODELLER http://salilab.org/modeller/ [10], YASARA http://www.yasara.org/) by supplying the raw amino acid sequences in fasta format. The generated results have to be interpreted carefully to find the structural changes in the mutant protein. However such interpretation and analysis on the molecular dynamics is not straightforward and simple.

We have derived a simple algorithm called modict to predict the effect of mutations on the structure of the protein. It is complementary to the protein modeling tools mentioned above, as it requires the 3D protein structures predicted by these tools. The algorithm takes into account the global structural changes in the 3D protein model. These structural changes are measured in means of the change in **R**oot **M**ean **S**quare **D**eviation (Δrmsd) and the corresponding residue number in the protein sequence.

## 2 Methods

### 2.1 Algorithm

Let *A*_*i*_ denote the rmsd value of a given amino acid at *i*^*th*^ position resulting from comparison of two models in a cartesian space defined by *V*(*i*, *A*_*i*_). Assuming the entire length of a protein with N residues is 1 unit, then the unit area of the rectangle enclosed by two consecutive amino acids can be approximated by:

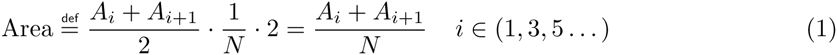

If a given domain is enclosed by *i*^*th*^ and *j*^*th*^ amino acid residues then the area spanned by the domain can be expressed as:

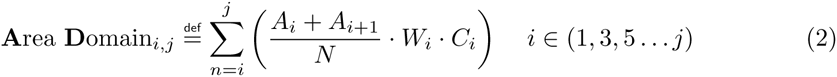

where *W*_*i*_ and *C*_*i*_ denote optional weight and conservation scores respectively which are usually provided by the training and iteration modules (users can attain as well). Of course the aforementioned area does not solely result from the mutation. An error value can be expressed in terms of overall rmsd (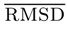; generated by SWISS-model):

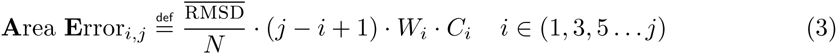

A total area can be defined from equations 2 and 3 (ad=Area Domain, ae=Area Error):

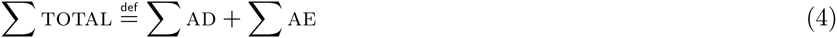

Above formula is a generalization for multiple domains. In case there is only one domain between residues *i* and *j*, than the total area simply is AD_*i*, *j*_ + AE_*i*, *j*_. A raw score (Γ) can be expressed in terms of:

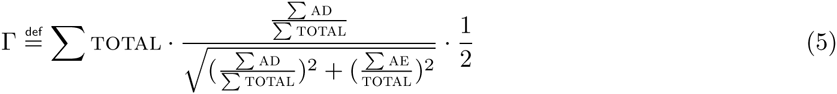

It is noteworthy that for a given interval, *AD* and *AE* are not guaranteed to be equal, even if the regions taken into consideration spans the entire protein. While *AD* is obtained from per residue rmsd, *AE* is obtained from 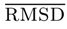. *AD/TOTAL* and *AE/TOTAL* should be considered as 2 orthogonal vectors. modict is designed to work with specific protein domains where i and j designate the start and end of a domain. For modict to perform optimal, it is important that the domains which are most critical for the functionality of the protein are chosen. This can be literature findings or can be predicted by the iteration script which is included in the software package (see section 2.3).

The difference (*δ*) between equations 2 and 3 is important to discern background signal from actual effect:

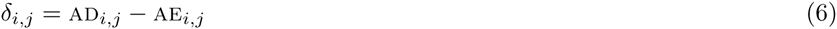

The significance (*γ*) of the difference depends on the length of the domain and the standard deviation of the individual rmsd values:

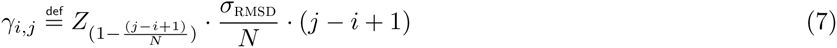

where *Z*_*x*_ denotes the Z score of (100 · *x*)^*th*^ percentile and *σ* denotes the standard deviation. Assuming that the rmsd values are distributed in a Gaussian distribution, the Z-score derived significance score gives an idea about how much of the domain residues account for the large rmsd values. From equations 6 and 7, a coefficient of significance (*κ*) can be defined:

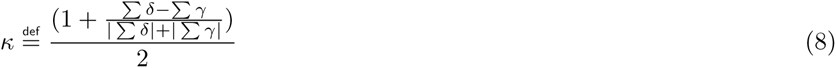

In the equation 8 above Σ*δ* or Σ*γ* denotes the total sum of *δ* or *γ* between all specified domain intervals such as *δ*_*i*, *j*_ + *δ*_*m*, *n*_ + *δ*_*u*, *w*_…. Equations 5 and 8 can be combined to express a final score:

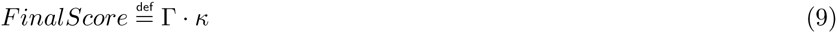

The criteria of evaluating the score can be performed via 2 different approaches as outlined in sections 2.2 and S1.2. In a fraction of cases, comparison of modict scores requires calculating thresholds and these thresholds are calculated via a *K* parameter. Beware that this is not the same coefficient as in equation 8. This parameter is a measure of the highest p-value attainable with a given accuracy. The *K* parameter is calculated from known list of mutations listed in table S1. For more information for the usage of this parameter refer to section S1.2.

### 2.2 modict methodology

The algorithm of modict is based on rmsd values of superimposed wildtype and mutant proteins. For calculating, rmsd values, a 3D protein model is required of both the wildtype and mutant case, which is calculated by using the i-tasser and phyre2 servers. After construction of the 3D models, the generated pdb files are used as input for a script included in modict which will extract the necessary rmsd values. For the purpose of testing modict, amino acid sequence of wildtype and mutant renin, Tubb2b, Btd and Smpdl proteins (uniprot id: P00797, Q9BVA1, P43251, P17405) were submitted to the automated i-tasser and phyre2 servers. PAH and ACADM (tables 1,2) were submitted to the automated phyre2 server. For further details on specific settings, see section S1.1. modict can be supplied with optional weight (min:0,default:10) and conservation(min:0,max:11,default:1) scores which are both array vectors (single number per line in a text file). Multiplying all entries of the weight and conservation file by a constant does not change the result. Both files are optional and not mandatory for modict to work. However, they can be used to give higher priority to certain regions. The default set up attains 1 to both conservation and weight scores.

**Table 1.** Mutations in PAH.

| Mutation | Residual Activity | Score |
| --- | --- | --- |
| Y414C | 28 | 0.112 |
| R241C | 25 | 0.136 |
| A403V | 32 | 0.125 |
| R261Q | 30 | 0.071 |
| E390G | 75 | 0.086 |
| R68S | 98 | 0.157 |
| I65T | 29 | 0.153 |
| V245A | 50 | 0.126 |
| L48S | 39 | 0.247 |
| F39L | 96 | 0.136 |
| D415N | 72 | 0.072 |
| A395P | 15 | 0.139 |
| A104D | 26 | 0.091 |
| R408Q | 55 | 0.063 |
| P211T | 72 | 0.185 |
| V388M | 43 | 0.15 |
| R241H | 23 | 0.131 |
| I306V | 39 | 0.161 |

Mutations in PAH with their residual enzyme activity and MODICT scores are listed. MODICT scores are generated taking into account the catalytic domain (143-410; [34]).

**Table 2.**
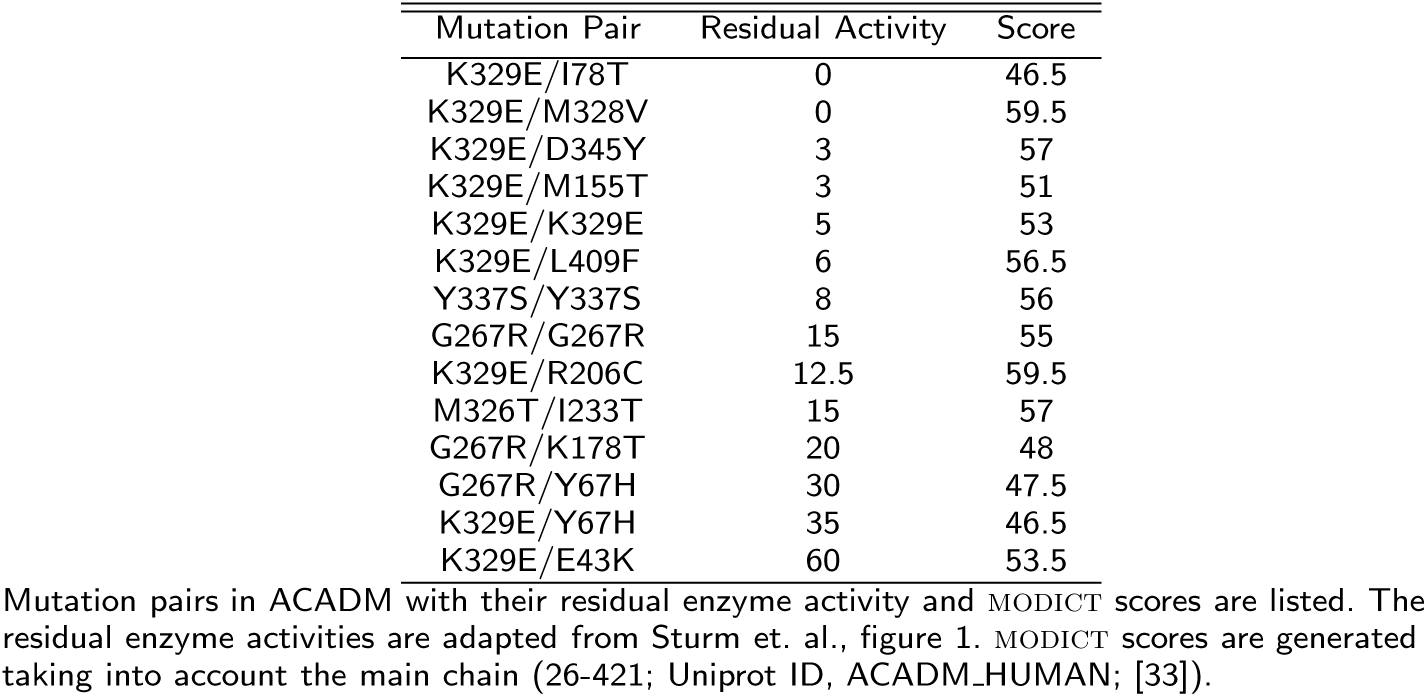
Mutations in ACADM.

Conservation scores are generated by aligning reviewed sequences of the protein of interest in different species from UniProt (http://www.uniprot.org/). It is a simple text file of one conservation score per line and generated using the jalview utility.

modict requires a user generated per-residue rmsd file as well. We have developed a script which can be supplied to swiss-pdb. This script extracts the rmsd values from superimposed WT (wildtype) and MT (mutated) ․pdb files to a file.

modict score interpretation makes use of a negative and positive control. As negative control, a superimposition between the wildtype protein and a refined model of the same wildtype protein (in some cases, a known benign mutation can also be used instead of refined wildtype, see sections 2.4 and S1.2). For the positive control, superimposition between the wildtype protein and a known pathogenic variant can be used. The scores for the negative and positive control can as such be used as a scale for the modict result of the protein variant of interest. A more mathematical approach to modict score interpretation is given in sections S1.2, 3.2, S1.3 and figure 7.

### 2.3 Training and Iteration

As will be described throughout the section 3, modict is designed to work with distinct domains which are critical for protein functionality. Often however, this information is not readily available. In order to meet these needs, modict comes with a training and iteration module where a random number approach is used to approximate a good candidate weight score combination as in figures 2, 4, 6, 8 and 9.

**Figure 1.**
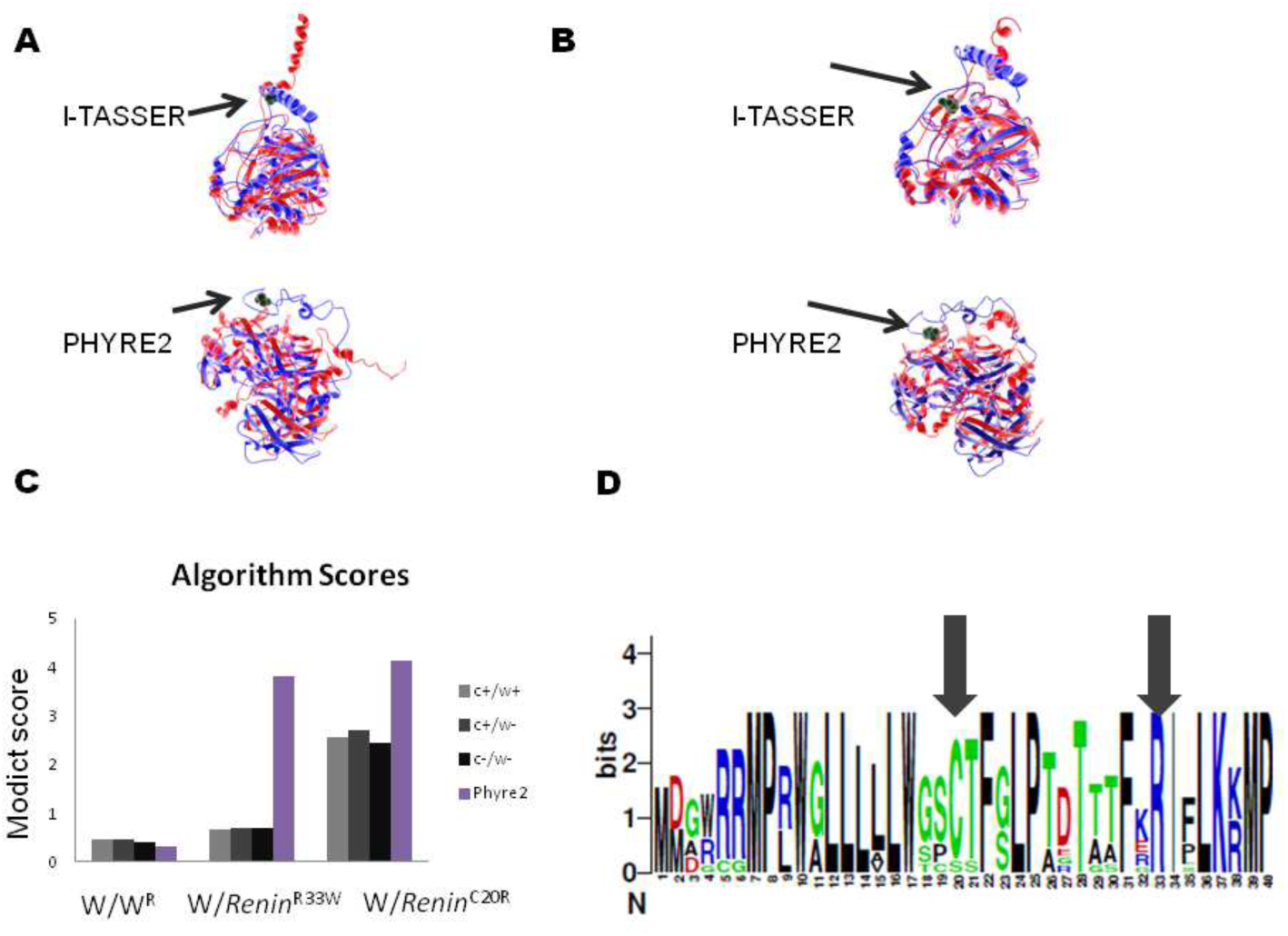
3D models of wildtype and mutated Renin. **A**. Wildtype (blue) and *Ren*^*p.C*20*R*^ (red) models are superimposed with the cysteine residue (green, Van der Waals) marked with arrow. Models generated with different modeling algorithms are indicated. **B**. Another variant in the signal sequence, *Ren*^*p.R*33*W*^ (red) does not result in a change to the same extent as. The wildtype arginine residue (green, Van der Waals) is marked with arrow. Graphical representation of algorithm scores, **C**. Absolute values of modict scores obtained from pairs; negative control (left, light gray; score: 0.455), wildtype against *Ren*^*p.R*33*W*^ (middle, light gray; score: 0.670) and positive control (right, light gray; score: 2.570). Algorithm scores with or without conservation (c) and weight (w) scores are also indicated (dark gray, black, see table S1). For comparison, algorithm scores generated using models from phyre2 is also indicated. Like black bars, these are raw modict scores generated without conservation and weight parameters. **Sequence logo of the renin signal peptide. D.** Residues 1-40 of reviewed renin sequences in UniProt database have been aligned. Note that both R33 and C20 are highly conserved, however algorithm scores significantly differ in case of i-tasser. modict scores were generated taking into account the main chain (residues 67-406, uniprot, P00797). (*W* = wildtype, *W*^*R*^ = refined wildtype)

**Figure 2.**
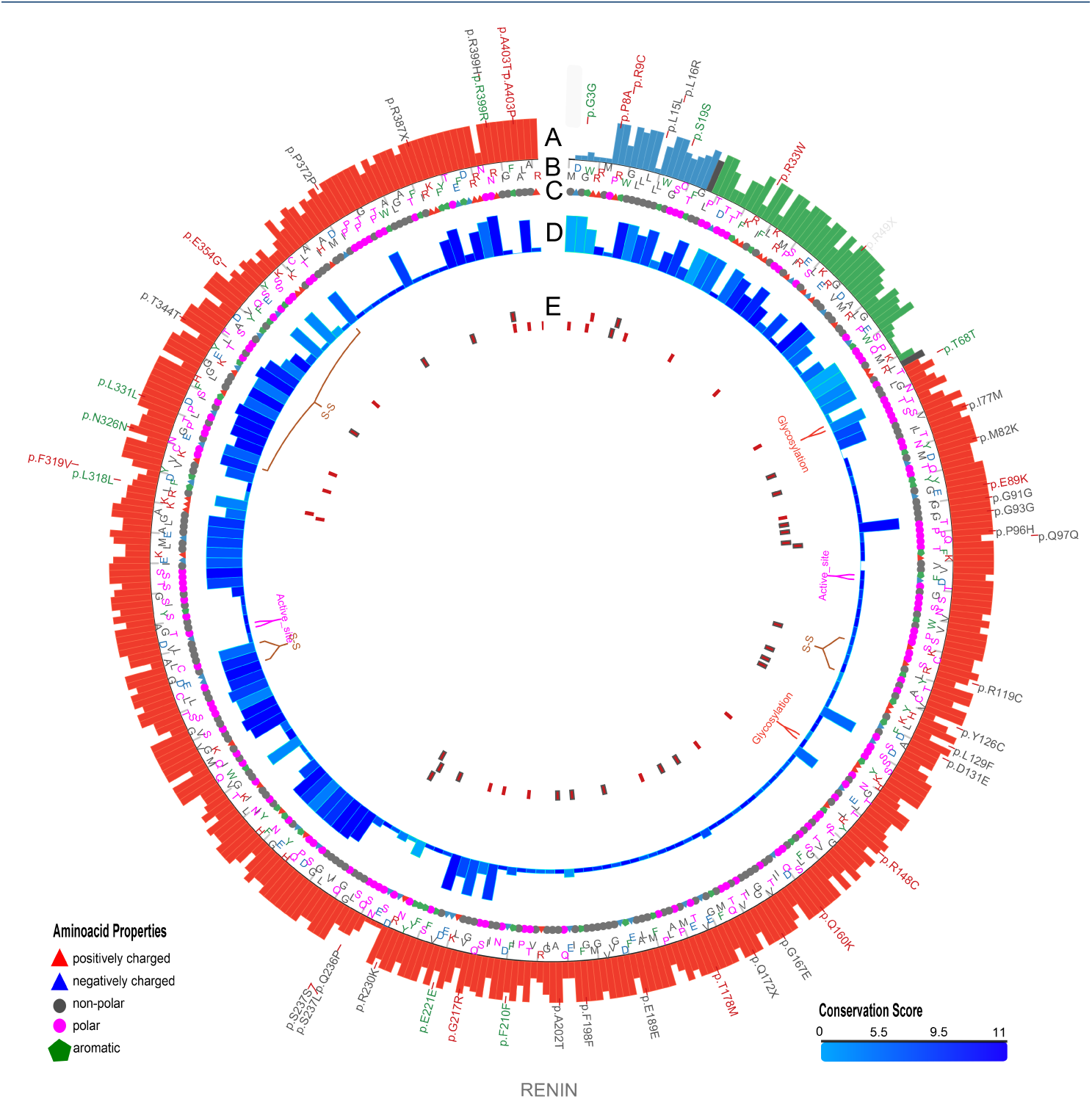
Plot showing conformational differences in *renin*^*C*20*R*^. Outermost layer indicates reported SNVs (Single Nucleotide Variants; gray, not validated; red, non-synonymous; green, synonymous) from dbSNP 138. **A.** Conservation scores represented as a histogram (blue, signal peptide; green, propeptide; red, domain). These values are generated as described in section 2.2 and are not related to modict score. **B** and **C.** Amino acid sequences with residues colored according to their property (positively charged, red triangle; negatively charged, blue triangle, non-polar, gray circle; polar, pink circle; aromatic ring, green hexagon). **D.** Iterative modict scores of individual residue pairs (algorithm, Eq.1) resulting from comparison with *renin*^*WW*^ and *renin*^*R*33*W*^. Each blue histogram bin designates the contribution of a residue pair to the overall modict score (Higher bars mean more contribution as well as more the adverse effect of that residue pair on structural stability). These histogram bins are generated by iterative modict algorithm and are colored according to conservation. **E.** Important regions, SNVs and Indels (insertion-deletions) are marked with boxes. Red boxes represent SNVs whereas pink boxes represent Indels. Gray bordered boxes represent unvalidated changes. (S-S = disulphide bond)

**Figure 3.**
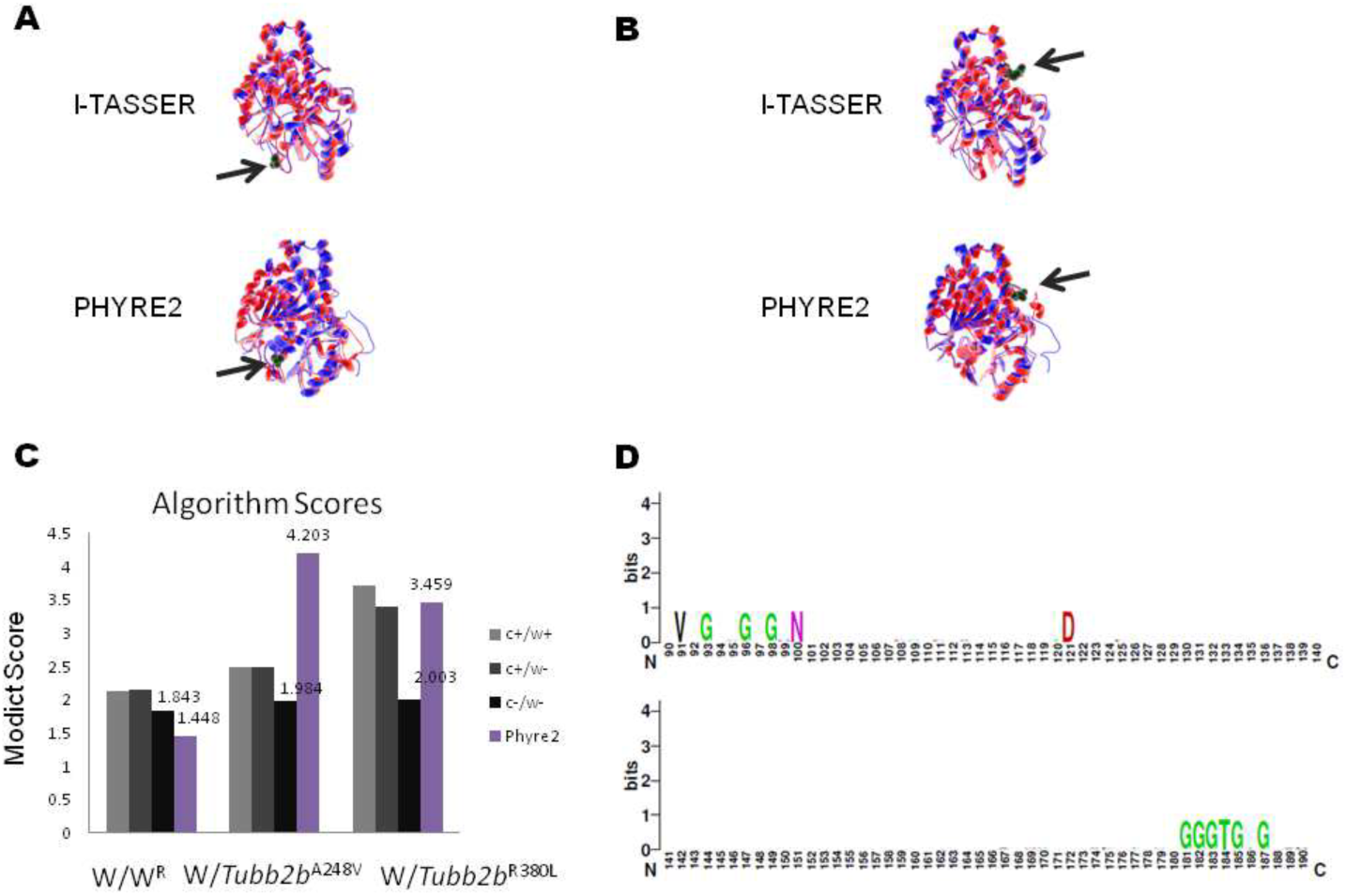
3D models of wildtype and mutated tubulin molecules, **A**. Superimposition of wildtype (blue) and *Tubb2b*^*p.A*248*V*^ (red) models. The alanine residue is rendered with Van der Waals radii (green, gray arrows). Models generated with different modeling algorithms are indicated. **B**. Structural comparison between wildtype (blue) and *Tubb2b*^*p.R*380*L*^ (red) models. The arginine residue rendered with Van der Waals radii (green, gray arrows). **Graphical representation of algorithm scores. C.** Absolute values of algorithm scores obtained from pairs; negative control (left, light gray; score: 2.129), wildtype against *Tubb2b*^*p.A*248*V*^ (middle, light gray; score: 2.485) and wildtype against *Tubb2b*^*p.R*380*L*^ (right, light gray; score: 3.721). For comparison, algorithm scores generated using models from FHYRE2 is also indicated. Like black bars, these are raw modict scores generated without conservation and weight parameters. **D. Sequence logo of conserved Tubb2b regions**. Residues 91-100 and 139-144 of Tubb2b have been conserved since their divergence from the FtsZ proteins. Consequently, during algorithm calculations they have received a weight score of 20 instead of default value. Scores with/without conservation or weight attributes are indicated in C. modict scores were generated taking into account the entire backbone (residues 1-445, uniprot, Q9BVA1). (*W* = wildtype, *W*^*R*^ = refined wildtype, c = conservation, w = weight score)

**Figure 4.**
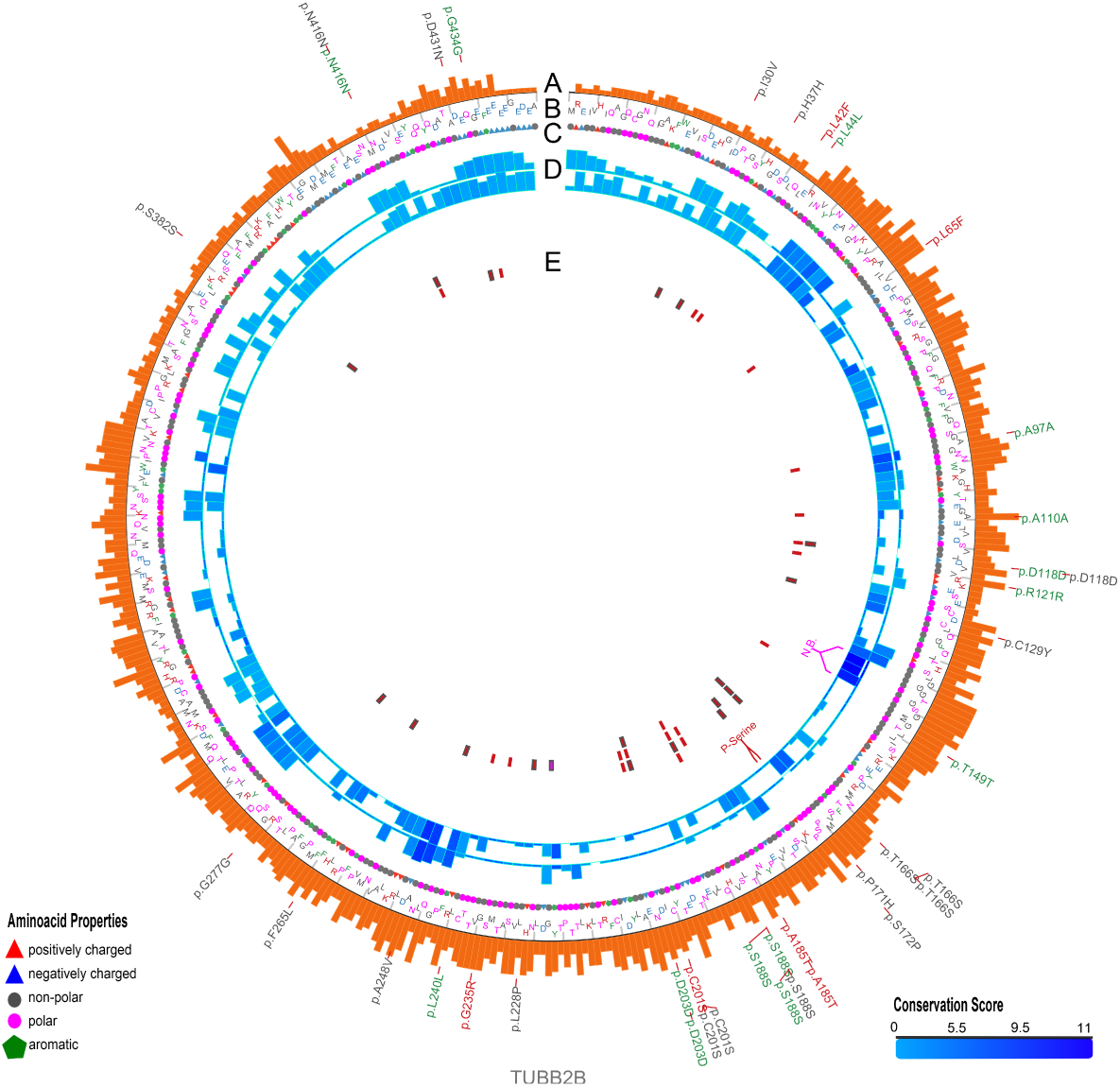
Plot showing conformational differences in *Tubb2b*^*A*248*V*^ and *Tubb2b*^*R*380*L*^. Outermost layer indicates reported SNVs (gray, not validated; red, non-synonymous; green, synonymous) from dbSNP. **A**. Conservation scores represented as a histogram. These values are generated as described in section 2.2 and are not related to modict score. **B** and **C**. Amino acid sequences with residues colored according to their property (positively charged, red triangle; negatively charged, blue triangle, non-polar, gray circle; polar, pink circle; aromatic ring, green hexagon). **D**. Iterative modict scores of individual residue pairs (algorithm, Eq.1) resulting from comparison with *Tubb2b*^*WT*^. Top layer belongs to *Tubb2b*^*A*248*V*^ whereas bottom layer belongs to *Tubb2b*^*R*380*L*^. Each blue histogram bin designates the contribution of a residue pair to the overall modict score (Higher bars mean more contribution as well as more the adverse effect of that residue pair on structural stability). These histogram bins are generated by iterative modict algorithm and are colored according to conservation. **E**. Important regions, SNVs and Indels are marked with boxes.

**Figure 5.**
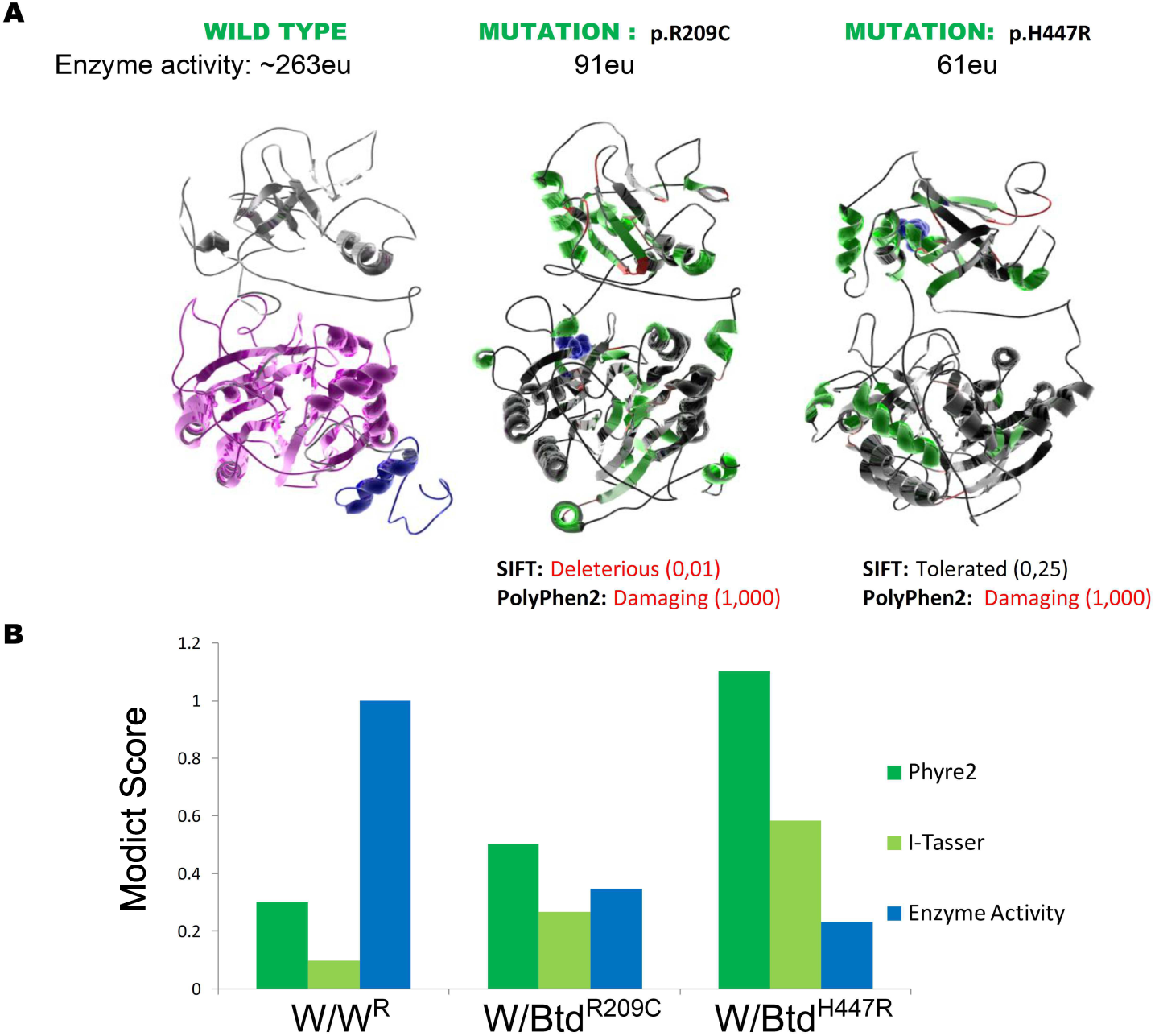
3D models of wildtype and mutated biotinidase. **A**. 3D biotinidase model generated by i-tasser (A, left). Pink residues (57-363) designate the CN-Hydrolase domain whereas the blue residues (1-41) designate the signal peptide. **Effect of p.R209C and p.H447R mutations on protein structure (A, middle, right)**. *Btd*^*WT*^ (left) is compared to p.R209C (middle) and p.H447R (right) in means of changes in secondary structure (no change, black; helix to strand, light green; strand to helix, dark green; helix to coil, light red; strand to coil, dark red; coil to strand or helix, green). The mutated R209 and H447 residues are depicted with blue Van Der Waals radii and their polyphen2/sift scores and residual enzyme activity are indicated. **Comparison of modict scores and residual enzyme activity, B**. modict scores from models generated by i-tasser (negative control, 0.096; p.R209C, 0.266; p.H447R, 0.584) and phyre2 (negative control, 0.301; p.R209C, 0.504; p.H447R, 1.102) were compared with experimentally measured enzyme activity (wildtype 263eu, p.R209C, 91eu, p.H447R, 61eu) scaled to 1. Ratios of modict scores and [1/enzyme activity] are in concordance with each other. (*W* = wildtype, *W*^*R*^ = refined wildtype)

**Figure 6.**
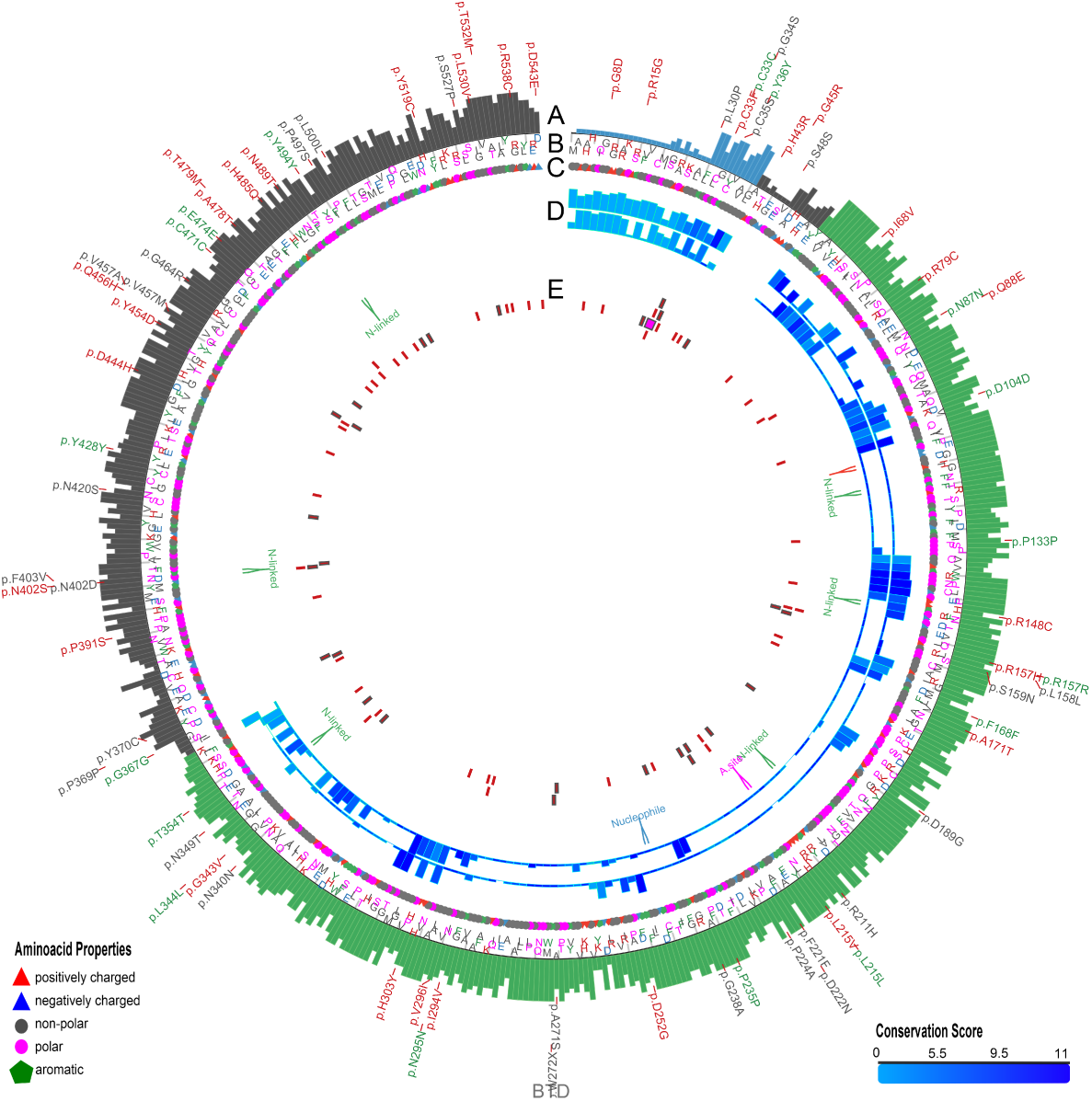
Plot showing conformational differences in *Btd*^*R*209*C*^ and *Btd*^*H*447*R*^. Outermost layer indicates reported SNVs (gray, not validated; red, non-synonymous; green, synonymous) from dbSNP. **A.** Conservation scores represented as a histogram (blue, signal peptide; green, CN-hyrolase domain). These values are generated as described in section 2.2 and are not related to modict score. **B** and **C.** Amino acid sequences with residues colored according to their property (positively charged, red triangle; negatively charged, blue triangle, non-polar, gray circle; polar, pink circle; aromatic ring, green hexagon). **D.** Iterative modict scores of individual residue pairs (algorithm, Eq.1) resulting from comparison with *Btd*^*WT*^. Top layer belongs to *Btd*^*R*209*C*^ whereas bottom layer belongs to *Btd*^*H*447*R*^. Each blue histogram bin designates the contribution of a residue pair to the overall modict score (Higher bars mean more contribution as well as more the adverse effect of that residue pair on structural stability). These histogram bins are generated by iterative modict algorithm and are colored according to conservation. Only scores belonging to domain regions re shown. **E.** Important regions, SNVs and Indels are marked with boxes. (A.site = active site)

**Figure 7.**
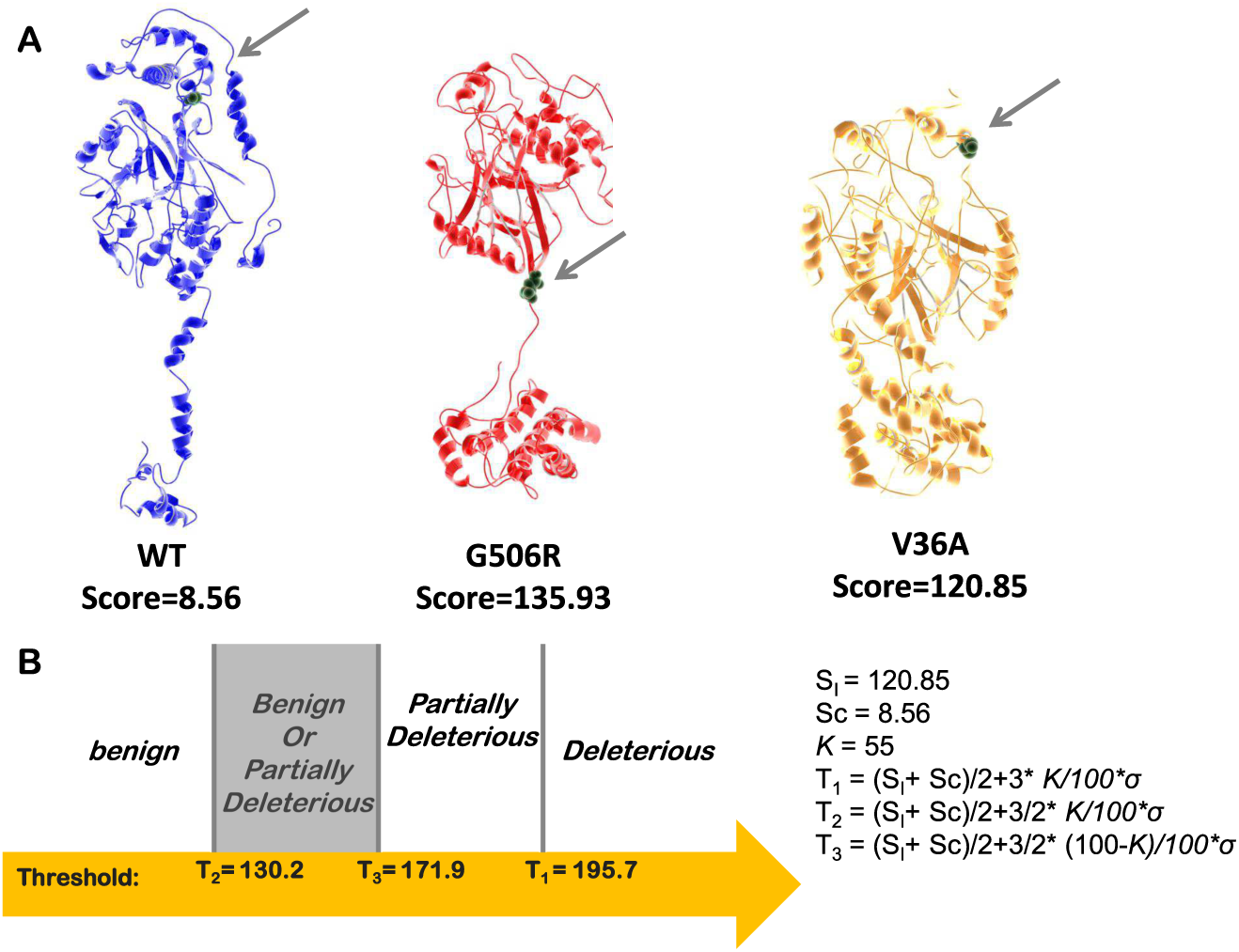
Classification of Smpd1^G506R^. **A**. Wildtype (blue), *Smpd1*^*G*506*R*^ (red) and *Smpd1*^*V*36*A*^ (orange) models are shown. The original position of glycine in wildtype, the substitution site in *Smpd1*^*G*506*R*^ and the alanine 36 in *Smpd1*^*V*36*A*^ are marked with gray arrows. Models have been further refined using the modrefiner. A negative control score was generated by superimposing the refined wildtype on the initial wildtype whereas a known benign score was generated by superimposing the refined *Smpd1*^*V*36*A*^ on the initial wildtype. A score for the test mutation was generated in the same manner. modict scores were generated taking into account the entire backbone (residues 1-629). **B**. Thresholds were calculated as shown in the right and the G506R mutation was classified based on the calculated score bracket as shown in the left. The value of kappa can be updated using the roc.pl script.(*σ* = *standard deviation of S*_*I*_ *and S*_*C*_)

**Figure 8.**
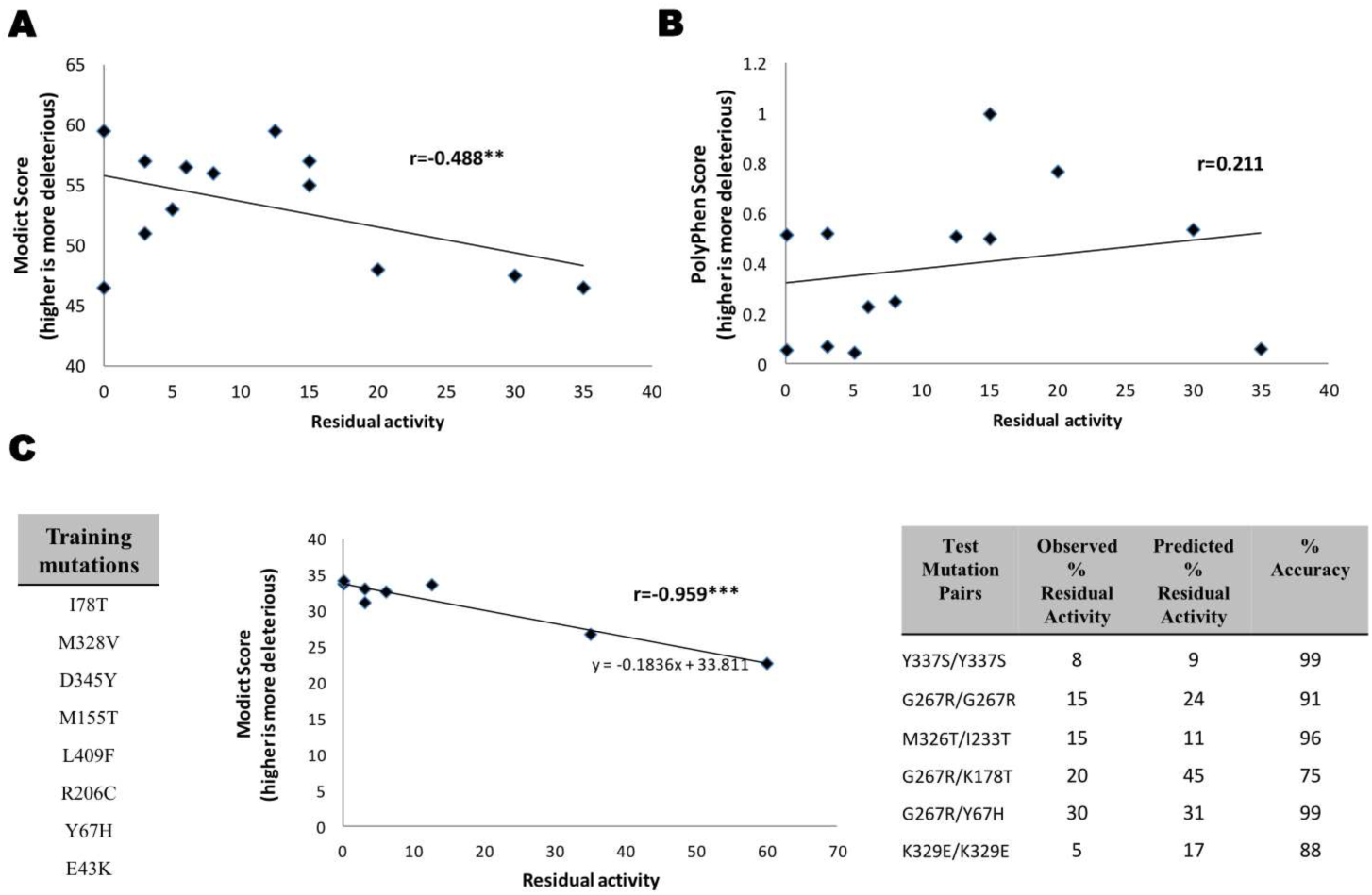
modict scores of ACADM mutations. **A**. Mutation pairs were plotted based on their enzymatic activity and the average of their modict scores. modict scores or residual activities that are 2 standard deviations away from the data average was excluded which corresponded to exclusion of only 1 data point (residual activity 60, modict score 53.5). The remaining data points had a correlation coefficient of −0.488 with a p-value of 0.044 according to 1 tailed t-distribution. **B**. Same mutations were plotted with polyphen2 scores instead which yielded a positive correlation coefficient of 0.211 with p-value of 0.244. **C**. 8 out of 14 mutation pairs in table 2 harbored a p.K329E variant where homozygotes for this mutation only had 5 percent of wildtype activity. Assuming significant portion of residual activity coming from the other variants, these 8 variants (lower left) were used as a training dataset for modict. After training, modict was able to find a weight score combination with a correlation coefficient of −0.959 (lower mid). Using the trendline obtained by least squares method, the residual activity of 6 other mutation pairs (that did not include the trained mutations) were guessed. modict was able to achieve 91 percent accuracy (lower right).(** = *p* < 0.05; *** = *p* < 0.001)

**Figure 9.**
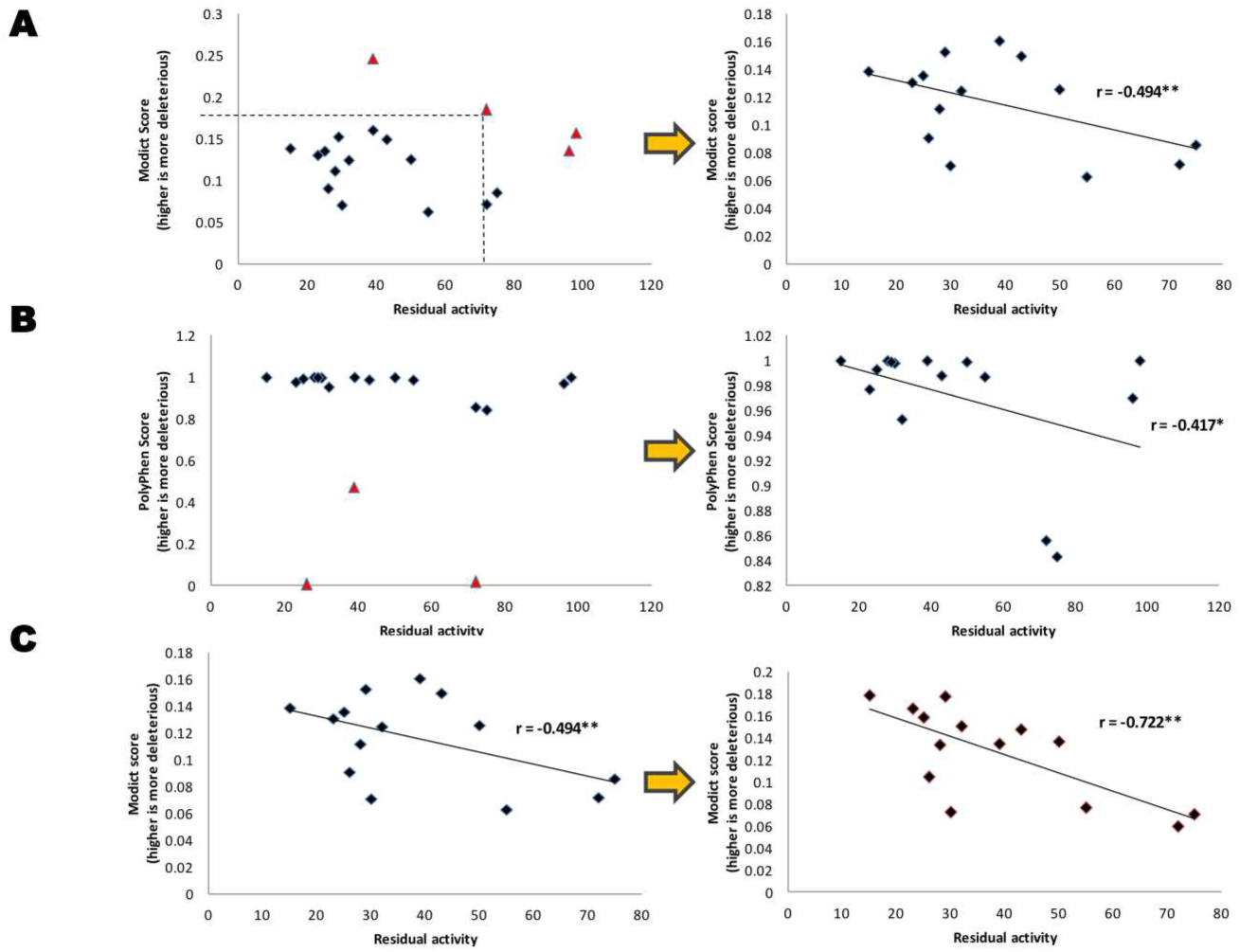
modict scores for partially deleterious PAH mutations. **Top Left**. Mutations with residual activity in PAH with their respective modict scores are plotted. Triangles indicate data points that are 2 standard deviations apart from the mean (both residual activity and modict score) of rectangle data points. **Top Right**. Outliers that are two standard deviations apart from the mean are removed and the correlation coefficient is calculated. modict scores are negatively correlated with residual activity (r=-0.494). The exact p-value of the correlation coefficient is 0.036 based on 1-tailed t-distribution. **Middle Left**. The same comparison was applied to polyphen2 scores. Triangle data points indicate the outliers. **Middle Right**. Likewise, polyphen2 scores were negatively correlated with residual activity (r=-0.417). However, the exact p-value of the correlation coefficient was 0.062 based on 1-tailed t-distribution. **Lower Left**. The training module of modict were used on the same mutations. **Lower Right**. The training module of modict was able to achieve a weight score configuration that yielded a more significant p-value of 0.002. (* = *p* < 0.1; ** = p < 0.05)

The training module accepts a list of paired modict scores and enzymatic activity (or any measure of residual protein function that is determined experimentally). It tries to find an optimal weight score combination for each residue that yields the highest possible Pearson’s correlation (one would expect enzymatic activity and modict scores to be negatively correlated). The user has control over the iteration process by regulating several parameters such as the number of rounds to iterate. Even then, improvement of initial correlation varies from protein to protein and depends on the number of mutations to be trained with.

modict package also comes with an iterator module to identify regions of a protein that contribute the most to the overall modict score (figures 2, 4 and 6). The iteration algorithm automatically attains weight scores between 0 and 10 to residues: the higher the weight score, the more the contribution of that residue pair to the overall modict score. modict uses a random number approach to approximate a significant combination. Although the computation process can be cumbersome under certain conditions, current approach performs well with comparison of many models simultaneously. Such an example is given in figure 10 where mutations that preserve more than or equal to 50 percent of residual activity are compared to two relatively more severe mutations.

**Figure 10.**
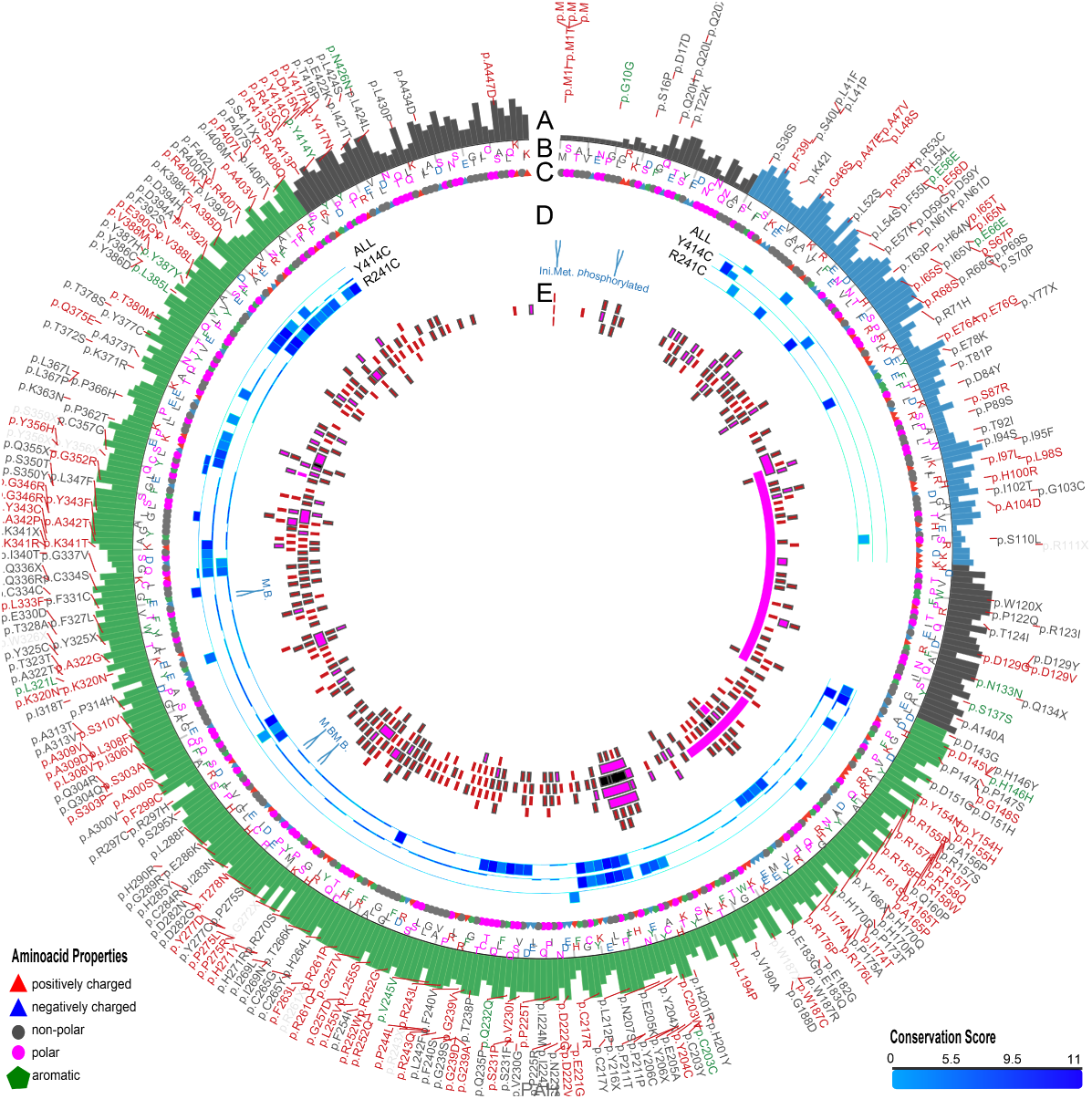
Plot showing conformational differences in *PAH*^*E*390*G*^, *PAH*^*V*245*A*^, *PAH*^*D*415*N*^, *PAH*^*R*408*Q*^, *PAH*^*Y*414*C*^, *PAH*^*R*241*C*^. Outermost layer indicates reported SNVs (gray, not validated; red, non-synonymous; green, synonymous) from dbSNP. **A.** Conservation scores represented as a histogram (blue, ACT domain; green, catalytic domain). These values are generated as described in section 2.2 and are not related to modict score. **B** and **C.** Amino acid sequences with residues colored according to their property (positively charged, red triangle; negatively charged, blue triangle, non-polar, gray circle; polar, pink circle; aromatic ring, green hexagon). **D.** Iterative modict scores of individual residue pairs (algorithm, Eq.1) resulting from comparison of mutations with residual enzyme activity less than 50 percent (more severe) against mutations with residual activity greater than 50 percent (less severe, table 1). Each blue histogram bin designates the contribution of a residue pair to the overall modict score (Higher bars mean more contribution as well as more the adverse effect of that residue pair on structural stability). These histogram bins are generated by iterative modict algorithm and are colored according to conservation. Single residue pairs with high blue bars are much less significant than consecutive “blocks” of high blue bars. Scarcity of these blocks in topmost layer (label: all) points to the fact that different regions are affected in each mutation. *PAH*^*Y*414*C*^ and *PAH*^*R*352*C*^ are compared to less severe mutations individually (middle and bottom layers). Note the differences in regions that are affected the most in each mutation. **E**. Important regions, SNVs and Indels are marked with boxes.

When the iteration algorithm of modict is used, it generates an automatic and interactable output as shown in figure 11. The user can choose to display amino acids with certain properties or just visualize the change in regions that correspond to a domain. The user may wish to know if residues with high modict score are also conserved which can be seen from the color coding. For a more comprehensive explanation of how to interpret iterator results please refer to modict documentation.

**Figure 11.**
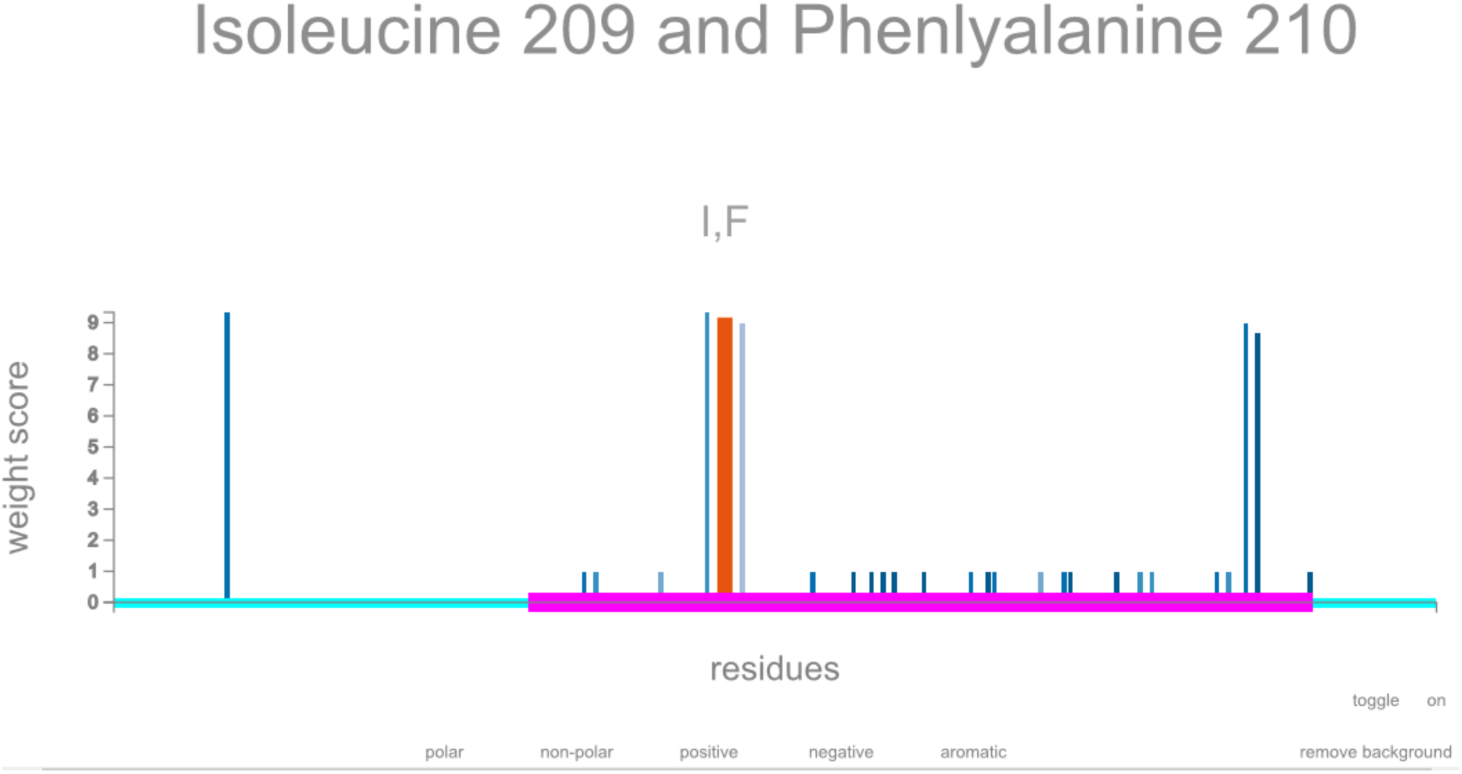
Automatically generated interactable output of iterative modict scores. Individual modict scores of residue pairs are plotted along the protein with an interactable interface. Annotation data is automatically stored with the use of modict. Histograms are automatically colored according to conservation data. Amino acids with different properties can be displayed separately. Pink regions highlights the functional domain. Data is taken from comparison of *PAH*^*Y*414*C*^ against *PAH*^*E*390*G*^, *PAH*^*V*415*N*^ and *PAH*^*R*408*Q*^. Only the amino acids with aromatic ring is displayed. Mouse over amino acids (209 I and 210 F) are highlighted. For a more comprehensive explanation of how to interpret iterator results please refer to modict documentation.

### 2.4 roc curve generation

One of the challenges to construct a receiver operating characteristic curve (roc) for an algorithm that generates a continuous range of output rather than a qualitative output (deleterious or benign) is to build a parametric classification system. This can be achieved by recalculating thresholds for a given set of mutations with known outcome while varying the levels of stringency (a measure of how rigorous the thresholds are constructed). Subsequently, this can be plotted against the p-value (a measure of how correctly the mutations are classified) In principle, mutations are not only completely benign or deleterious but spread through a range of variable residual protein activity/function. In addition to a negative control which is usually Δrmsd between wildtype and a refined wildtype model or wildtype and a benign model, another score from Δrmsd between wildtype and a given benign/deleterious/partial model should be used. This allows the user to construct a hypothetical distribution of scores and thus determine the likelihood of a test score being benign, deleterious or partial. Such a script is included in the modict package. The user can import his calculated scores from new models and update the current RGC plot shown in figure 12. Data used to generate the plot is listed in table S1.

**Figure 12.**
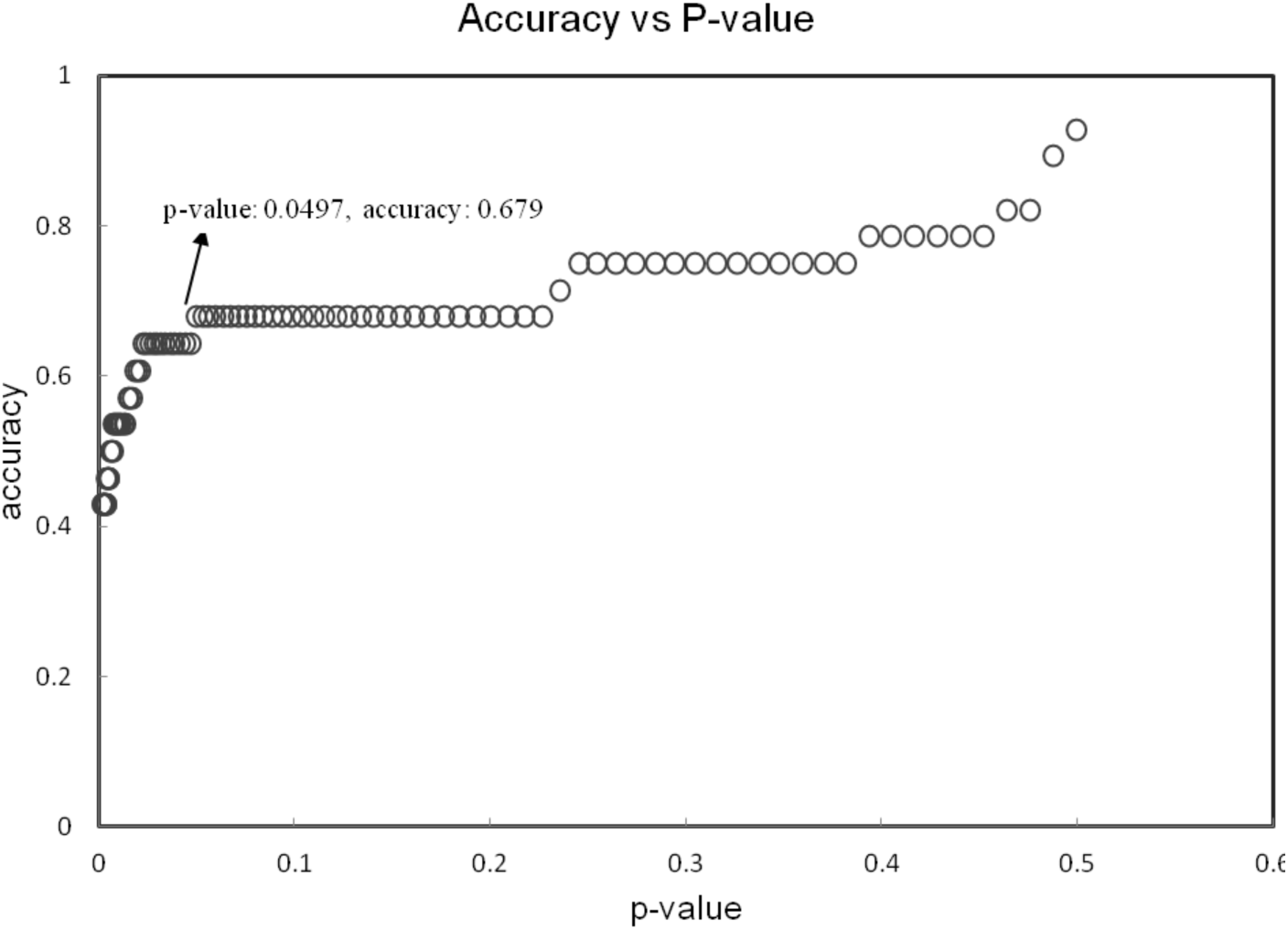
roc curve. Trio groups (negative control, test, positive control) are tested for decreasing levels of stringency measured as a parameter depending on the standard deviation of the negative controls and the positive controls. There is a trade off between the p-value and the stringency. As stringency decreases, accuracy increases, however the increase in accuracy can be explained progressively less by the measurements of the algorithm (increasing p-value and decreasing significance). The data used to generate the above plot is indicated in table S1. The script for generating the data above is included in modict package.

### 2.5 Output

modict, supplied with the rmsd file, gives as an output an algorithm score, which is a float value without units.

## 3 Results

We have derived a simple algorithm modict to predict whether a mutation is deleterious or not based on the rmsd obtained from superimposed mutated and wildtype 3D structures. The 3D protein structures in this study were modeled by i-tasser and phyre2, however other modeling algorithms can be used as well. The mathematical model underlying modict can also incorporate the information from conservation and weight scores. An iteration algorithm to determine the regions that account the most for the calculated score is also available with modict. modict is not only a prediction tool, but also a tool to scrutinize changes in the protein structure independent of the score.

The algorithm was tested on 6 different proteins which belong to different protein families. The chosen mutations were of different nature in order to minimize bias. modict scores were interpreted by two methods,either correlating them with experimental metrics like enzymatic activities, or using the scores for ordinal classification (deleterious, benign, partially deleterious etc.). The first method requires modict scores for at least 3 mutations with experimentally verified enzyme activities for predicting the effect of unknown mutation. Then, the modict scores and the enzymatic activity of the known mutations are plotted in a scatter plot and a trend-line is set by the least squares method. By observing the trend-line the enzymatic activity of your mutation of interest can be traced. The advantage of this approach is the ability to use the training module on modict for a subset (or the entire set) of mutations to increase the initial Pearson’s r correlation coefficient. This method was applied on Btd, Pah and Acadm mutations (see tables 1,2 and figure 3.3).

The second method is used when there are less than or equal to 2 mutations. However a negative control modict score is required for comparison. This method was applied on Renin, Tubb2b and Smpdl mutations (see sections 3.1,3.2 and 3.4). Regardless of the method, higher modict scores mean more deleterious.

Throughout this paper modict scores have both been used as ordinal classifiers (benign, partially deleterious, deleterious etc.) and continuous variables to measure correlation. In all of the tested cases in this study whether conservation scores and/or weight scores were used or not is indicated. Concerning the examples given in this article, modict performs better without conservation scores.

Throughout the results section, output of the iteration algorithm (residues that contribute the most to a modict score) was represented using I-PV as shown in figs 2,4,6 and 10 [11].

### 3.1 Renin p.R33W

Renin is one of the main components that regulates the main arterial blood pressure via the renin-angiotensin system and is initially secreted as a propeptide with a 67 amino acid long signal sequence [12]. Mature renin does not have this signal sequence and is 37kDa long [13]. A novel heterozygous mutation c.58T>C (p.C20R) was found in all affected members of a family with autosomal dominant inheritance of anemia, polyuria, hyperuricemia and chronic kidney disease [14].

Another variant p.R33W suspected to be benign resides within the same signal sequence (http://www.ncbi.nlm.nih.gov/projects/SNP/snp_ref.cgi?rs=11571098;-http://web.expasy.org/variant_pages/VAR_020375.html). Several prediction algorithms were tested on this variant previously [15]. In this example, conservation scores generated by multiple sequence alignment of reviewed Ren (renin) sequences were also used by the algorithm as an additional factor (section S1.3). Based on domain annotations, residues that are involved in various interactions were also given a weight score of 20 instead of default value (10, section S1.3). Figure 1C and figure 2 show the algorithm results associated with these mutations.

We also provided wildtype and mutated Renin fasta files to automated FHYRE2 server and received models for the same variants. Wildtype Renin score was 0.328 whereas p.R33W and p.C20R scores were 3.816 and 4.128 respectively. Based on these scores p.R33W variant should be classified as deleterious. As mentioned previously, the p.R33W is of unknown significance due to its low frequency (dbSNP, <1%). Although a study has claimed that it significantly reduces Renin biosynthesis (http://www.ashg.org/2014meeting/abstracts/fulltext/f140120880.htm), to our knowledge it has not yet been published. The Renin example demonstrates that modict scores are not totally independent from the models provided to it. For more detailed explanation for using modict scores as an ordinal classifier, please refer to the manual and section S1.3.

### 3.2 Tubb2b p.A248V and p.R380L

Tubulins are the main components of microtubules on which dynein and kinesin motor proteins bind. Together with intermediate filaments and microfilaments, they form the cytoskeleton which plays a major role in intercellular trafficking, cell-cell interactions, junctions and cellular migration [16]. Tubulins are ubiquitously expressed in all human tissues. However mutations in these proteins mostly affect tissue types that rely on their functionality the most during development such as cells of neuronal or glial origin [17,18]. Almost all mutations in tubulins result in Malformations of Cortical Development (MCD) [19]. Mutations in *TUBB2B* result in polymicrogyria spectrum of malformations. [20–26]. 2 *de novo* mutations in Tubb2b, namely p.A248V and p.R380L in 2 unrelated patients of Turkish and Belgian origin and 1 patient of French-Canadian origin respectively were identified and tested for their modict scores [21].

Figure 3 (C) and figure 4 show the algorithm results associated with these mutations. Scores without weight and conservation parameters (section S1.4) for wildtype, *Tubb2b*^*p.A248V*^ and *Tubh2h*^*p.R380L*^ were 1.843, 1.984 and 2.003 respectively. Choosing the wildtype as control (*S*_*C*_) and *Tubb2b*^*p.R380L*^ as known deleterious mutation (*S*_*K*_), the threshold *T*_1_ was calculated as 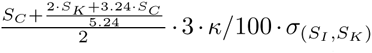.

The value for *T*_1_ was 1.945 which was lower than the *Tubb2b*^*p.A248V*^ score (*σ* = standard deviation, *κ* = 55). This means that the *Tubb2b*^*p.A248V*^ mutation is indeed deleterious.

Wildtype and mutated fasta files were provided to the automated phyre2 server. modict scores in the absence of weight and conservation parameters for wildtype, *Tubb2b*^*p.A248V*^ and *Tubb2b*^*p.R380L*^ were 1.448, 4.203 and 3.459 respectively. Choosing *Tubb2b*^*p.A248V*^ as the known deleterious variant, the *T*_1_ threshold is 3.200 which is lower than the *Tubb2b*^*p.R380L*^ score. As a result, modict scores generated by both i-tasser and phyre2 models agree on the nature of the variants.

### 3.3 Btd p.H447R and p.R209C

Biotinidase is an enzyme that is encoded by the *BTD* gene. Low enzyme activity interferes with the cycling of biotin and if left untreated, it may lead to neurological and cutaneous issues [27]. In this example, a case with experimentally verified results from 2 patients will be used and compared with modict scores [28]. The genotype of the patients in the aforementioned study were c.1330G>C (p.D444H)/c.1340A>G (p.H447R)[patient 1] and c.557G>A (p.C186Y)/c.625C>T (p.R209C)[patient 2]. Both former mutations (c.1330G>C in patient 1 and c.557G>A in patient 2) were null mutations meaning that the experimentally measured residual enzyme activity belongs to the latter mutations [27,28]. The residual enzyme activity in the patients were 61eu (enzyme units) and 91eu respectively (population mean 263*eu*). modict scores were generated using 2 different modeling algorithms (i-tasser, phyre2) and results were compared with residual enzyme activity as shown in figure 5 [8,29]. Conservation scores were generated by aligning reviewed biotinidase sequences from UniProt (*Homo sapiens, Rattus norvegicus, Mus musculus, Bos taurus, Takifugu rubripes*) by using Clustal Omega (http://www.ebi.ac.uk/Tools/msa/clustalo/) and the resulting scores (min, 0; max, 11) corresponding to 1-543 residues of Btd were given to modict [30]. Supplying or not supplying the conservation scores do not significantly alter the *score*_modict_/*enyzmatic* – *activity* ratios as can be seen from table S1.

The modict scores were generated by taking into account functionally important regions (residues 57-363, 402-403 and 489-490; uniprot, P43251). These functionally important regions can generally be found in UNIPROT. As seen in figure 5, both PHYRE2 and i-tasser scores are proportional to corresponding enzymatic activities. Although there are only 2 mutations, taken together with the negative control score, raw modict scores without any conservation or weight files correlate strongly with enzymatic activity (phyre2: *r* = –0.805; i-tasser: *r* = –0.838).

### 3.4 Mutations in Sphingomyelin phosphodiesterase-1

Sphingomyelin phosphodiesterase-1 is an enzyme (Uniprot ID: ASM_HUMAN) located in lysosomes and responsible for conversion of sphingomyelin to ceramide. Deficits in enzyme activity or reduction in the enzyme concentration result in an inborn error of metabolism grouped under the name Niemann-Pick disease (type A and B) [31]. Several polymorphisms exist that are frequent amongst control populations. One example of such variant is the p.V36A located in the signal sequence. Another variant that is often mistaken as deleterious is p.G506R [32]. Using FHYRE2 to model wildtype, figure 7 demonstrates the procedure of classifying the p.G506R mutation. Since the known p.V36A variant is benign (with a score of *S*_*K*_), the *S*_*I*_ score is substituted directly by *S*_*K*_. Based on the calculated thresholds, the p.G506R mutation was correctly classified as “partially deleterious or benign”. The procedure to use modict as an ordinal classifier using thresholds is further elaborated in the manual and in the discussion section.

### 3.5 Mutations in Medium Chain Acyl-CoA Dehydrogenase

Medium chain acyl-coa dehydrogenase (MCAD, Uniprot ID: P11310, NP_000007.1) is an enzyme encoded by the *ACADM* gene. MCAD deficiency is one of the most common deficits in mitochondrial *β*-oxidation. MCAD is the enzyme responsible for breaking down medium-chain fatty acids. Deleterious mutations that reduce the enzyme activity result in clinical symptoms such as hypoglycemia, hepatic and neuronal dysfunction [33]. Enzymatic activity data of homozygous/compound heterozygous patients carrying 2 deleterious mutations have been adapted from Sturm *et al.* as shown in table 2 [33]. Mutated proteins were modeled using phyre2 and superimposed on wildtype MCAD which was generated by submitting wildtype fasta file to the phyre2 server. For each mutation pair the modict score was the average of the modict score of individual mutations (direct summation without average only expands the graph on one axis). Rather than using modict as a classifier, the main goal was to see if the modict scores correlates with the real experimental measurements. modict scores correlated negatively with the enzymatic activities as shown in figure 8.

Because higher modict scores denote more deleterious effect, as the residual activity increases, it’s well expected for modict scores to go down which results in negative correlation. As shown in figure 8, the initial Pearson’s correlation coefficient was −0.488. Although not very strong, it is important to underscore that modict is the first attempt to achieve such degree of correlation between prediction and experimental outcome from user generated 3D protein models. Figure 8 also compares correlation of polyphen2 scores with enzymatic activity which did not yield significant concordance with experimental results.

Figure 8 also depicts the use of the training module of modict. Table 2 lists the compound heterozygous mutations used for correlations in figure 8. Eight of the mutation pairs in table 2 share a near-null deleterious p.K329E mutation where homozygotes for this variant has five percent residual activity. Thus, we have trained modict with these eight mutations and then used the trendline (calculated by least squares method) to guess the enzymatic activity of other remaining mutation pairs in table 2. As shown in figure 8 (lower right), modict was able to achieve 91 percent accuracy. The MCAD example demonstrates the possibility of developing an enzyme specific panel without the need of very large datasets for training of modict.

### 3.6 Mutations in *PAH*

The last example is about pheynlketonurea (PKU), an enzymatic defect that manifests itself with the deficiency in phenylalanine hydroxylase (PAH), a phenylalanine to tyrosine converter with the aid of tetrahydrobiopterin (BH4). It is an autosomal recessive disease with both copies of *PAH* carrying deleterious mutations. The ample decrease in PAH activity results in elevated phenylalanine blood concentration. If the elevated phenylalanine concentration is left untreated, it can lead to mental retardation with structural brain changes visible on a MRI. Deleterious mutations in *PAH* affects variably the level of enzymatic activity. Data regarding such mutations can be found in several studies [34,35]. Comparison of the generated modict scores after excluding outliers shows that the scores of individual mutations were negatively correlated with residual enzyme activities as shown in figure 9 (Pearson’s r = −0.494). Similarly, polyphen2 scores correlated negatively with experimental measurements but to a lesser degree (Pearson’s r = −0.417). Using the training module for the 14 mutations in figure 9 further improved the initial correlation coefficient from −0.494 to −0.722.

## 4 Availability and Future Directions

### Discussion

modict is an algorithm which predicts whether a mutation is deleterious or not. This is based on the rmsd obtained from superimposing mutated and wildtype 3D protein structures. Modeling was done here by using i-tasser and phyre2, although alternatives can be used as well. The mathematical model underlying modict can also incorporate the information from conservation and weight scores. An iteration algorithm to determine the regions that account the most for the calculated score is also available with the package.

There are two ways to make use of modict scores. The first way is to convert the scores into an ordinal classification system, which requires a negative control. The second way is to correlate experimental results with modict scores as shown in the *BTD, MCAD* and *PAH* examples. The bottleneck in this approach is to find several known mutations in the protein of interest with available enzymatic activities or an equivalent measurement. However, this method allows an extrapolation between modict scores and residual protein activity. By using the MODICT training module, one can further optimize the linear relationship between modict scores and residual enzyme activities. Although overall rmsd values and significance is taken into account by the algorithm, modict’s accuracy still depends on the models generated by the user. Unlike polyphen2 and sift, modict scores are not normalized and vary depending on the length of protein, rmsd values between residues, overall rmsd, regions that are taken into account etc. Therefore individual modict scores should not be seen as values indicative of deleterious or benign nature, but should always be interpreted in relation to their negative/positive controls or in relation to known enzyme activities.

### Reporting results with Modict

When reporting results using modict, users should provide the parameters they used together with the tool. Several of these parameters are key factors in reproducibility of the results. One of these parameters is the modeling algorithm used (phyre2, i-tasser etc.) and the sequence of the protein submitted to the server. The other parameter is the regions that are taken into account (residue numbers, domains etc.) when calculating the modict score. The user should also indicate the conservation and the weight scores used, if any. If the training algorithm is used, than the mutations used for training and the output weight score combination should be reported as well. If the user has followed the ordinal classification method, then she/he should also indicate how the negative control score was generated. Lastly, the users should also indicate the superimposition method used for generating the rmsd values. For example, superimposition based on alpha carbon has been used throughout this article.

### Limitations

modict is a tool that is not independent on the models generated by the modeling algorithm of choice. The Renin case is a good example for this where models generated by phyre2 and i-tasser gave different modict scores. Moreover, consistency in superimposition techniques used between models and the portion of the protein that is actually modeled (full length protein modeling is usually more reliable than partial modeling of distinct domains) significantly affect the outcome. Many modeling servers also include a confidence key together with the results which are useful to judge the quality of starting models. In general, since the wildtype model will be the main model where test and known mutated models are superimposed on, a low quality model will make it harder to discern between scores. Another issue is that many modeling servers have amino acid limits on submitted fasta files which are generally below 2000. This might make the evaluation of large proteins harder. As modeling algorithms advance, several of these issues will be resolved. Another drawback is that all structural deviations from a given wildtype model is perceived towards the deleterious spectrum whereas in reality there are also gain of function mutations. In that case, it is possible to modify the range of weight scores to include negative values as well.

### Future directions

It is important to underline that modict has no universal training dataset. This means that the algorithm itself (without any weight or conservation parameters) is able to reflect and capture portion of the physio-chemical interactions that determine the outcome of pathogenicity, at least for the proteins demonstrated in this article. In later stages the conservation scores or more importantly the weight scores can be used to train modict on a protein basis. For instance certain combinations of weight scores that yield a higher correlation coefficient for a given enzyme panel can be generated. We planning to train modict on variety of proteins and upload the trendlines for each modeling algorithm so the end user would only have to upload his/her mutation’s modict score without having to train the algorithm manually.

A systematic database of modict scores could be very beneficial for additional variant filtering in Next Generation Sequencing analysis as the utilization of protein structures files is not adequately implemented. We are planning to store user-submitted modict scores for this purpose. modict is a fully automated algorithm that comes with a variety of scripts to analyze the effects of mutations on protein structure. Unlike most other mutation predictors, modict uses. pdb files and can simultaneously compare multiple models for differences in topology. All the models used for this article can be downloaded together with the modict package from https://github.com/modict/modict.

## Acknowledgments

Ibrahim Tanyalcin received funding from Scientific Fund Willy Gepts and the Foundation Marguerite Delacroix. AJ received funding from the Research Foundation Flanders.

## Supplementary Section

### S1.1 3D protein models and annotation

Amino acid sequences of wildtype and mutant renin, Tubb2b, Btd and Smpd1 proteins (uniprot id: P00797, Q9BVA1, P43251, P17405) were submitted to the automated i-tasser and phyre2 servers. PAH and ACADM (tables 1,2) were submitted to the automated fhyre2 server with the intensive mode selected (including wildtype fasta files). The obtained 3D models of renin, Tubb2b, Btd and Smpd1 were energy minimized on deepview-swiss-pdbviewer (http://www.expasy.org/spdbv/ [9,10]) via 2 cycles of steepest descent consisting of 50 steps each and 1 cycle of conjugate gradient consisting of 200 steps with a minimum energy difference(ΔE) of 0.01kJ/mol together with a harmonic constraint of 100 kJ/mol. Models were further refined using modrefiner (http://zhanglab.ccmb.med.umich.edu/ModRefiner, [36]). For each query a trio pair was constructed by comparing the ratio of the final scores between wildtype/wildtype-refined, wildtype/test and wildtype/mutated where the first and last components serve as negative and positive controls respectively. Images were post-processed with pov-ray v3.6 (http://www.povray.org). Models can be downloaded together with the modict package. The annotation of mutations in this article is in concordance with the Human Genome Variation Society (HGVS, http://www.hgvs.org/).

### S1.2 Using modict scores

There are two ways to make use of modict scores. The first way is to convert the scores into an ordinal classification system, which requires a negative control. A negative control score is made by superimposing WT-refined over WT pair (see section S1.1). For the first way, the negative control score can be generated by resubmitting your wildtype model to a refinement server such as modrefiner. Or the user can use his own *in-silico* pipeline for model refinement. After refinement, the user should superimpose the refined wildtype model on the wildtype one to generate a negative control score. The important point here is to apply the same refinement procedure to mutated models before superimposing them on the wildtype. However, to justify the use of this system, the user has to have only 2 mutations (one with known effect) with no enzymatic activities to correlate with. The reason is, if there are multiple known mutations, then there will be multiple thresholds. The second approach yields higher resolution and alleviates the problem of multiple thresholds. Supposing you have 3 modict scores (negative control: *S*_*C*_, test: *S*_*T*_, any score from known mutation: *S*_*K*_), it is possible that your known mutation might be deleterious, partially deleterious or benign. The first two cases requires you to reverse calculate an hypothetical benign (*S*_*I*_) such that 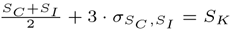 (*σ*_*x*, *y*_ = standard deviation of x and y) for a deleterious 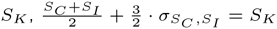 for a partially deleterious *S*_*K*_ and simply *S*_*I*_ = *S*_*K*_ for a benign *S*_*K*_ Than the critical value can be expressed as 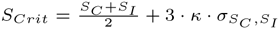. is greater than *S*_*Crit*_, than the query mutation is classified as deleterious. If the difference 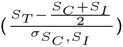 between 3*κ* and 1.5*κ* standard deviations than the mutation is classified as partially deleterious and finally if the difference is less than 1.5*κ*, than the mutation is classified as benign. The value of *κ* is determined from the roc (receiver operating characteristic; refer to section 2.4) plot with the data listed in table S1 and the current value is 0.55.

The second way is to correlate experimental results with modict scores as shown in BTD, MCAD and PAH (see sections 3.3, 3.5 and 3.6) examples. The bottleneck in this approach is to find several mutations in the protein of interest with available enzymatic activities or an equivalent measures.

### S1.3 Renin p.R33W

Conservation scores were generated by multiple sequence alignment of reviewed Ren (renin) sequences (Uniprot Entry names: reni1_mouse (*Mus musculus*), reni_canfa (*Canis familiaris*), reni_macmu (*Macaca mulatta*), reni_sheep (*Ovis aries*), reni2_mouse (*Mus musculus*), reni_human (*Homo sapiens*), reni_pantr (*Pan troglodytes*), reni_calja (*Callithrix jacchus*), reni_macfa (*Macaca fascicularis*), reni_rat *Rattus norvegicus*). Domain annotation was based on databases of prosite (http://prosite.expasy.org/), interpro (http://www.ebi.ac.uk/interpro/) [37, 38] and UniProt.

Using modict as an ordinal classifier requires calculating thresholds. Figure 1 scores are given for algorithm results generated taking into account weight and conservation scores. To focus on results generated solely by modict, scores generated without weight or conservation scores will be used which are indicated in table S1 and as black bars in figure 1C. To calculate thresholds, a *κ* value is also necessary which is generated based on the examples in table S1. Current value of *κ* is 55 (based on the mutations tested in this article). Users can update table S1 with additional data. In principle, more data points (mutations with known effect) will output a more realistic *κ* value. Taking the negative control score (*S*_*C*_) as 0.396, the known mutation score (*S*_*K*_) as 2.491, an imaginary benign score (*S*_*I*_) is calculated as 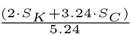. Next, the *T*_1_ threshold is calculated as 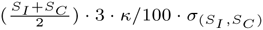 which in this case is 1.705 (*σ* = standard deviation). If your test score is larger than this value, then your mutation is classified as deleterious. The value of p.R33W (0.684) is smaller than this value which requires calculation of threshold *T*_2_ given by 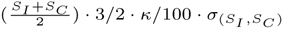. The value of *T*_2_ for this case is 1.247 which the p.R33W score is below and thus classified as benign. If the score would be larger than *T*_2_ but below *T*_1_, the variant would be considered as partially deleterious. Values of *κ* below 66 also enable calculation of *T*_3_ threshold which divides partially deleterious mutations into 2 classes: partially deleterious or benign and partially deleterious. For this example *T*_3_ calculation is not necessary.

### S1.4 Tubb2b p.A248V and p.R380L

Conservation scores were generated by aligning reviewed Tubb2b (*Homo sapiens, Mus musculus, Rattus norvegicus, Bos taurus, Xenopus laevis*), Tuba1a (*Homo sapiens, Mus musculus, Rattus norvegicus, Sus scrofa, Pan troglodytes, Cricetulus griseus*), Tubb3 (*Homo sapiens, Mus musculus, Rattus norvegicus, Bos taurus, Macaca fascicularis, Arabidopsis thaliana*) and FtsZ (*M. jannaschii, S. aureus, E. coli*) sequences from UniProt. Moreover, weight scores were attained based on alignment of FtsZ (*M. jannaschii, S. aureus, E. coli*) sequences with Tubb2b as shown in figure 3 (D).

### S1.5 Additional comments

As shown in figures 1 and 3, it is relatively clear to classify Renin^R33W^ compared to Renin^C20R^, however differences in the tubulin dataset are relatively small and thus calculation of score brackets is necessary. As a general rule of thumb, proteins that are evolutionarily conserved across species are more sensitive to missense mutations and this fact is reflected on the data by exhibition of closer modict scores between different mutations. This phenomenon can be observed by elevation of negative control scores like in figure 3 for the Tubb2b protein.

As previoulsy stated, there are two ways to make use of modict scores. The first way is to convert the scores into an ordinal classification system, which requires a negative control. The second way is to correlate experimental results with modict scores as shown in BTD, MCAD and PAH examples. The bottleneck in this approach is to find several known mutations in the protein of interest with available enzymatic activities or an equivalent measurements. The advantage of this method is to be able to omit the negative control score as the linear trendline (assessed by least squares method) becomes the main means of calculating predicted enzymatic activities. Another advantage is to be able to use the training module for modict. Training modict on subset of mutations increase the linear relationship between residual enzyme activities and modict scores. Consequently the new trendline can be used to remap enzymatic activities of new mutations as shown in MCAD example, figure 8.

modict should be seen as a tool rather than an “all in one” program to predict a variant’s pathogenicity. It is an attempt to standardize the usage of user generated 3D models for predicting the effect of mutations. modict is licensed under GPL and is composed of 7 scripts and 2 modules which ultimately aim to relate extracted rmsd values from mutated proteins with experimental results. Although overall rmsd values and significance is taken into account by the algorithm, modict’s accuracy still depends on the models generated by the user. Unlike polyphen2 and sift, modict scores are not normalized and vary depending on the length of protein, rmsd values between residues, overall rmsd, regions that are taken into account etc. Therefore individual modict scores should not be seen as values indicative of deleterious or benign nature; modict scores are unit-less. Rather than a universal threshold, the relationship between modict scores are important in their interpretation. The two methodologies for interpretation (ordinal classification and correlation) have been shown throughout this article. Comparison of modict scores are always done within the same protein. Therefore using large number of mutations from different family of proteins for bench-marking is not relevant in case of modict as opposed to mainly sequence-based predictors like polyphen2 and sift. This does not mean that information in sequence is obsolete, on the contrary, it means that modict allows users to approach the prediction process from a different angle.

**Table S1.** roc curve data.

| Wildtype | Given | Test | *Condition<sup>test</sup>* | *Condition<sup>given</sup>* | Conservation | Algorithm | Protein | *Mutation<sup>given</sup>* | *Mutation<sup>test</sup>* |
| --- | --- | --- | --- | --- | --- | --- | --- | --- | --- |
| 0.467 | 0.704 | 2.696 | deleterious | benign | alignment | I-TASSER<sup>†</sup> | renin | R33W* | C20R |
| 0.467 | 2.696 | 0.704 | benign | deleterious | alignment | I-TASSER<sup>†</sup> | renin | C20R | R33W* |
| 0.396 | 0.684 | 2.453 | deleterious | benign | default | I-TASSER<sup>†</sup> | renin | R33W* | C20R |
| 0.396 | 2.453 | 0.684 | benign | deleterious | default | I-TASSER<sup>†</sup> | renin | C20R | R33W* |
| 2.158 | 2.491 | 3.401 | deleterious | deleterious | alignment | I-TASSER<sup>†</sup> | Tubb2b | A248V | R380L |
| 2.158 | 3.401 | 2.491 | deleterious | deleterious | alignment | I-TASSER<sup>†</sup> | Tubb2b | R380L | A248V |
| 1.843 | 1.984 | 2.003 | deleterious | deleterious | default | I-TASSER<sup>†</sup> | Tubb2b | A248V | R380L |
| 1.843 | 2.003 | 1.984 | deleterious | deleterious | default | I-TASSER<sup>†</sup> | Tubb2b | R380L | A248V |
| 0.092 | 0.267 | 0.619 | partial | partial | default | I-TASSER<sup>†</sup> | Btd | R209C | H447R |
| 0.092 | 0.619 | 0.267 | partial | partial | default | I-TASSER<sup>†</sup> | Btd | H447R | R209C |
| 0.1 | 0.272 | 0.599 | partial | partial | alignment | I-TASSER<sup>†</sup> | Btd | R209C | H447R |
| 0.1 | 0.599 | 0.272 | partial | partial | alignment | I-TASSER<sup>†</sup> | Btd | H447R | R209C |
| 6.2 | 160.269 | 162.143 | deleterious | benign | alignment | I-TASSER | tmem | A198V | G212V |
| 6.2 | 162.143 | 160.269 | benign | deleterious | alignment | I-TASSER | tmem | G212V | A198V |
| 2.919 | 67.783 | 68.283 | deleterious | benign | default | I-TASSER | tmem | A198V | G212V |
| 2.919 | 68.283 | 67.783 | benign | deleterious | default | I-TASSER | tmem | G212V | A198V |
| 0.489 | 2.176 | 2.775 | deleterious | deleterious | alignment | I-TASSER<sup>†</sup> | ACADM | E43K | K329E |
| 0.489 | 2.775 | 2.176 | deleterious | deleterious | alignment | I-TASSER<sup>†</sup> | ACADM | K329E | E43K |
| 0.467 | 2.147 | 2.605 | deleterious | deleterious | default | I-TASSER<sup>†</sup> | ACADM | E43K | K329E |
| 0.467 | 2.605 | 2.147 | deleterious | deleterious | default | I-TASSER<sup>†</sup> | ACADM | K329E | E43K |
| 0.33 | 1.127 | 0.514 | partial | partial | alignment | PHYRE2 | Btd | H447R | R209C |
| 0.33 | 0.514 | 1.127 | partial | partial | alignment | PHYRE2 | Btd | R209C | H447R |
| 0.325 | 1.175 | 0.562 | partial | partial | default | PHYRE2 | Btd | H447R | R209C |
| 0.325 | 0.562 | 1.175 | partial | partial | default | PHYRE2 | Btd | R209C | H447R |
| 14.127 | 217.87 | 307.33 | benign | benign | alignment | PHYRE2 | Smpd1 | V36A | G506R |
| 14.127 | 307.33 | 217.87 | benign | benign | alignment | PHYRE2 | Smpd1 | G506R | V36A |
| 8.56 | 120.85 | 135.93 | benign | benign | default | PHYRE2 | Smpd1 | V36A | G506R |
| 8.56 | 135.93 | 120.85 | benign | benign | default | PHYRE2 | Smpd1 | G506R | V36A |

Each line constitutes a trio composed of a negative control (wildtype), a positive control (given) and a test. The results of all mutations are previously known and are written under conditions column. Different modeling algorithms are used to minimize bias and they are indicated. Proteins functional in its entirety are tested along the protein backbone whereas proteins with known domain annotations are tested for specific regions. (\*= Clinical significance unknown;no study in favor of adverse functional affect has been published in a scientific journal during the time of this project. †= Also modeled with PHYRE2 as demonstrated in the results section.)

**Table S2.**
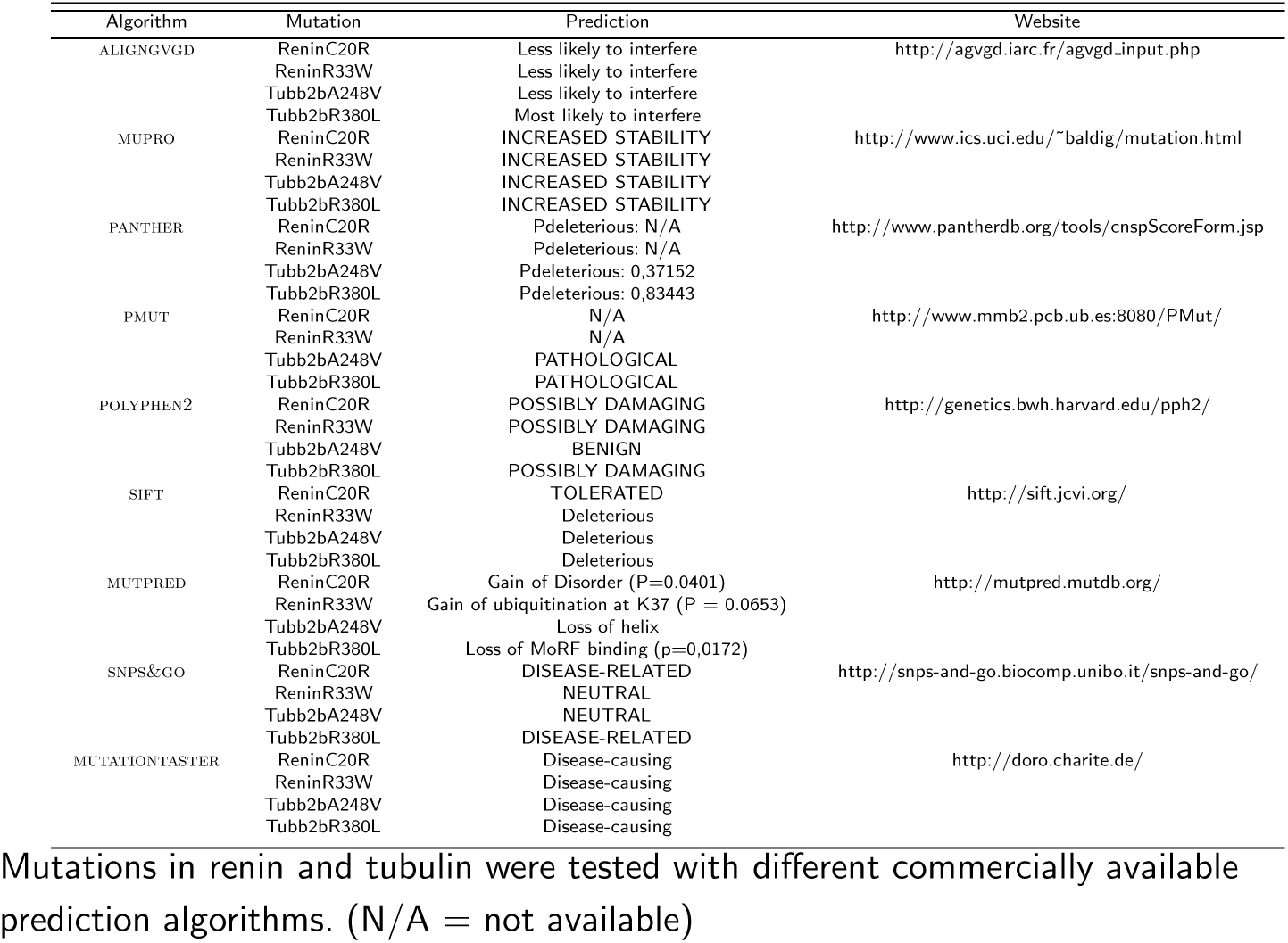
Mutations in renin and tubulin.

